# Modelling variability and heterogeneity of EMT scenarios highlights nuclear positioning and protrusions as main drivers of extrusion

**DOI:** 10.1101/2023.11.17.567510

**Authors:** Steffen Plunder, Cathy Danesin, Bruno Glise, Marina A. Ferreira, Sara Merino Aceituno, Eric Theveneau

## Abstract

Epithelial-Mesenchymal Transition (EMT) is a key process in physiological and pathological settings (i.e. development, fibrosis, cancer). EMT is often presented as a linear sequence of events including (i) disassembly of cell-cell junctions, (ii) loss of epithelial polarity and (iii) reorganization of the cytoskeleton leading to basal extrusion from the epithelium. Once out, cells can adopt a migratory phenotype with a front-rear polarity and may additionally become invasive. While this stereotyped sequence can occur, many in vivo observations have challenged this notion. It is now accepted that there are multiple EMT scenarios and that cell populations implementing EMT are often heterogeneous. However, the relative importance of each EMT step towards extrusion is unclear. Similarly, the overall impact of variability and heterogeneity on the efficiency and directionality of cell extrusion has not been assessed. Here we used computational modelling of a pseudostratified epithelium to model multiple EMT-like scenarios. We confronted these in silico data to the EMT occurring during neural crest delamination. Overall, our simulated and biological data point to a key role of nuclear positioning and protrusive activity to generate timely basal extrusion of cells and suggest a non-linear model of EMT allowing multiple scenarios to co-exist.

## Introduction

Epithelia are tight cell populations in which cells display an apicobasal polarity with apical cell-cell adhesions and basal cell-matrix adhesions. Epithelia can generate mesenchymal cells through a process called Epithelial-Mesenchymal Transition (EMT). EMT occurs iteratively during embryo development for morphogenesis and organogenesis^1^ but it is also involved in fibrosis and cancer^2^. In particular, EMT is critical for metastasis from carcinoma, the main cause of death in patients^3^. The main events occurring during EMT are a loss of cell-cell adhesion, a detachment from the extracellular matrix as well as changes in polarity and cytoskeleton dynamics. These steps are sometimes coupled with acquisition of invasive features such as expression of proteases^4^. Depending on the scale of the process (i.e number of cells involved), EMT can lead to the disassembly of the epithelium or basal extrusion of cells from the epithelium.

For communication and teaching purposes, EMT is often presented as a linear cascade controlled by an array of transcription factors implementing the molecular changes in a seemingly logical order. This theoretical sequence starts from the loss of cell-cell adhesion, followed by a loss of epithelial polarity, a breach in the local matrix and a subsequent extrusion of the cell body from the epithelium. Then, the mesenchymal cells, freed from their neighbors and the constraints of the epithelium, may adopt a migratory behavior depending on the local environment. This “engineering view” of the EMT process seems logical because detaching cells before starting migration is likely the most efficient option in terms of energy. However, this view is in striking contrast with what we know now about EMT in multiple systems.

EMT is no longer considered as a switch between opposite states but a progressive change of cell state that is rarely completed to a full mesenchymal phenotype and is reversible^2^. It allows cells to adopt hybrid/intermediate phenotypes along the E-M spectrum^5-7^. This means that, at the single cell level, there is variability with multiple scenarios that cells can use to go from E to M. While at the cell population level, there is heterogeneity with neighboring cells implementing different scenarios in parallel. This raises the question of the relative importance of the various events taking place during EMT and whether different scenarios are linked to various degrees of extrusion efficiency. Moreover, the effect of heterogeneity on the efficiency of extrusion is also unknown. Providing answers to these questions may be of clinical importance since phenotypic heterogeneity in tumors has been linked to therapeutic resistance in cancer and EMT was deemed one of the main drivers of this observed heterogeneity^8, 9^.

One of the best studied examples of physiological EMT is the basal extrusion of neural crest cells from the pseudostratified neuroepithelium during early embryonic development^10^. Neural crest cells are multipotent stem cells that form most of the peripheral nervous system, pigment cells, cartilages and bones of cephalic structures as well as smooth muscles, adipocytes, tendons or endocrine cells^11^. In amniotes (e.g. mammals, reptiles and birds), neural crest cells depart by performing basal extrusion towards the extracellular matrix before migrating. Neural crest cells express an array of transcription factors to implement EMT and launch cell migration. Interestingly, a wide diversity of gene expression profiles and cell behaviors has been observed in neural crest cells. In the anterior (cephalic) regions, hundreds of cells perform EMT in a short time window (6-10h)^12^, leaving the neural tube ***en masse***. By contrast, in posterior (truncal) regions, basal extrusion of cells from the neuroepithelium is spread out over several days^13^. In addition to variation in pace and cell numbers, heterogeneities in terms of gene expression have been observed. Known regulators of EMT in neural crest cells such as Ets1, Foxd3, Snail2, Zeb2 or Zic1 are not systematically co-expressed in all delaminating neural crest cells^12, 14-19^. Further, a diversity of strategies for cells to get out of the neuroepithelium has been documented^20, 21^. In these studies, in contrast with the “engineering view”, neural crest cells can translocate their cell body out of the epithelium without disassembling cell-cell adhesion and prior to a loss of epithelial polarity. In addition, protrusive activity has been seen before effective detachment from the epithelium and prior to the extrusion of the cell body^20, 21^.

Studying the variability and heterogeneity of EMT and their impact on basal extrusion ***in vivo*** would require being able to control adhesions and cell polarity independently, with single cell resolution and time control. Unfortunately, this is currently unachievable. One reason is technical. It is currently impossible to reliably perform, ***in vivo***, multiple gain/loss-of-function strategies with the required level of precision. Even if it was, a massive biological hurdle remains as none of the molecular effectors are specific to a given EMT step. Affecting cell-cell adhesion modulates polarity and vice versa^22, 23^. Similar feedbacks exist between cell-cell and cell-matrix adhesions^24^, including in neural crest cells^25^. Further, cytoskeleton dynamics are essential for localization of adhesion components but also for maintenance of polarity, protrusive activity as well as for cell division or interkinetic nuclear movements (INM). Thus, there are no neat experimental strategies to assess the relative roles of adhesion, polarity, protrusion, proliferation and INM during EMT ***in vivo***.

One way to circumvent such hurdles is to use computational modeling to be able to control each cell parameter independently in time and space. In the model, we can for instance impair cell-cell adhesion without affecting polarity or modulate independently cytoskeleton-related events such as apical constriction, INM or mitoses. So far, modelling has been instrumental in our conceptualization of EMT as a dynamic process in which cells toggle between E and M states, to bring forward the notion that EMT and its reverse process, Mesenchymal-Epithelial Transition (MET), are only partially symmetrical processes, to explore the link between E-M plasticity and stemness or to propose how non-genetic heterogeneity can emerge^8, 9, 26-28^. However, the relationship between gene expression of EMT regulators and implementation of specific steps such as cell junction disassembly or remodeling of polarity remains sketchy. Thus, such models do not allow to make prediction about the relative importance of EMT molecular and cellular events on extrusion efficiency. To tackle this problem, we built on a previously validated model of proliferating pseudostratified epithelium recapitulating the growth of the early neuroepithelium of the chicken embryo^29^ in which we now incorporate the time control of cell parameters (e.g. cell-cell adhesion, cell-matrix adhesion, INM) at single cell level. Using this model, we assessed the impact of epithelial destabilization on the ability of the cells to extrude apically or basally. We modelled (i) loss of cell-cell adhesion, (ii) detachment from the basement membrane and (iii) relaxation of apicobasal polarity. We implemented these various events with/without protrusive activity and with/without INM. We started by modelling the impact of a single event in an individual cell or homogeneous groups and progressively increased the complexity to model heterogeneous populations of cells performing various EMT scenarios in parallel.

Our simulations show that a multitude of EMT scenarios can lead to apical and basal extrusions from the epithelium with various efficiencies. The number of cells, as well as the timing and order of most events had only marginal effect on the relative efficiency of both types of extrusion. By contrast, we found that basal positioning of the nucleus at the onset of EMT and protrusive activity severely bias the timing and directionality of extrusion. Further, we show that heterogeneity acts as a destabilization factor boosting extrusions in both directions. According to these computational outputs, we reassessed some cellular and molecular aspects of EMT in chicken neural crest cells. Our biological results support the importance of basal positioning of nuclei and remodeling of ECM/cell adhesion in very early stages of EMT. In addition, we found a correlation between heterogeneity of neural crest cells and efficacy of extrusion ***in vivo***, supporting our conclusions from simulation studies.

Overall, our computational and experimental data lead us to propose that the cellular implementation of EMT is the result of an array of multiple inputs influencing the timing and directionality of extrusion. The observed diversity of cell behaviors in a cell population undergoing EMT may reflect the fact that one of the key step of basal extrusion (basal positioning of the nucleus) is not bound to a single cellular event. This reduces the pressure on the timing and order of events in EMT scenarios and may in itself lead to the emergence of a certain degree of heterogeneity. We generated a user-friendly online version of our model as a web-based, **<u>s</u>**tand-alone **<u>EMT</u>** simulat**<u>or</u>** (sEMTor, https://semtor.github.io/). Without any installation steps, anyone can run simulations corresponding to the initial conditions used in the various figures.

## Results

To simulate the various EMT-like scenarios, we generated a 2D agent-based model of proliferating pseudostratified epithelium (Figure 1a-b) derived from our previously established model^29^. Briefly, in the model, each cell is abstracted to a nucleus attached to a set of dynamic springs that represent the viscoelastic properties of the cell. These springs are terminated by an apical point ***a*** and a basal point ***b***. Apical points of adjacent cells are linked to one another by a contractile spring representing cell-cell adhesion. Basal point are attached to a simplified matrix represented by a straight non-deformable line. Basal points can only move along that line and cannot swap positions. Each nucleus (***N***) is made of two spheres: a hard core at the center and a soft core at the periphery. Hard cores cannot overlap while soft cores can but are subjected to a repulsion force. This allows to account for the deformation of nuclei that occurs at high density in pseudostratified epithelia without having to model actual changes in nuclear shape^29^. To maintain the stereotypical straight cell shape observed in pseudostratified epithelia, there is a straightness spring that imposes a flat 180° angle between the apical point, nucleus and basal point of each cell. Cells proliferate following a simplified cell cycle (Figure 1b) with passive springs in G1, S and early G2 phases. During late G2 and mitotic (M) phases the apical spring contracts to bring the nucleus on the apical side by generating pre-mitotic rapid apical movements (PRAM) driving interkinetic nuclear movements (INM) of nuclei. A complete description of the model is provided in Supplementary Information.

**Figure 1.**
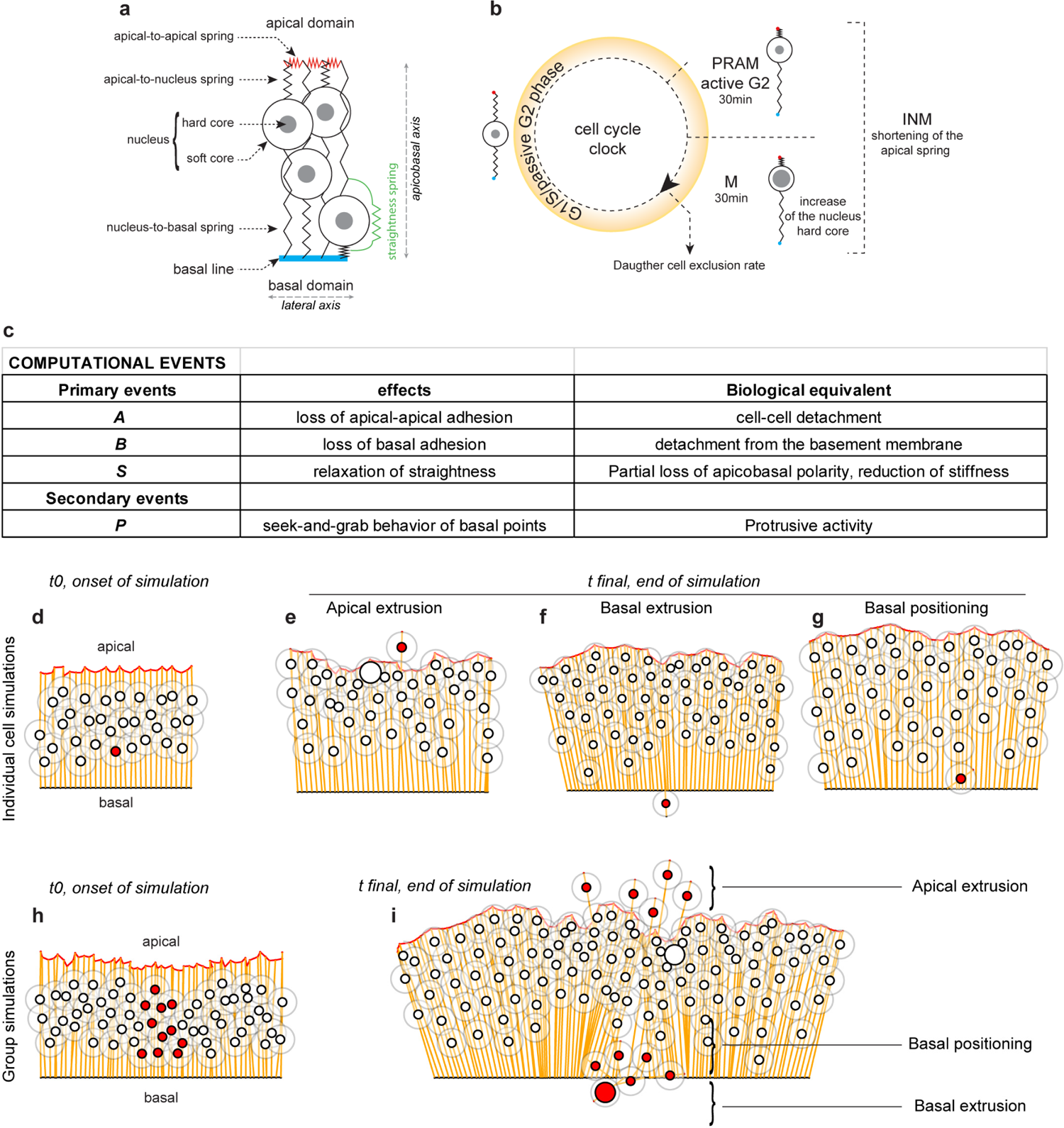
Presentation of the computational model a, Diagram representing the simulated cells with the various dynamic springs controlling cell-cell adhesion (apical-to-apical, red), cell-matrix adhesions (nucleus-to-basal spring, black), the viscoelastic-like properties of the cell body (apical-to-nucleus and nucleus-to-basal springs, black), the alignment of the apical point, nucleus and basal point (straightness spring, green). b, overview of the simplified cell cycle implemented in the model. The apical-to-nucleus spring contracts in active G2 and M phases, the hard core of the nucleus increases in M phase. c, Table of computational events used to simulate EMT-like scenarios throughout the study. Note that ***P*** is a secondary event as it can only occur if ***B*** happened. d, h, Initial organization of the tissue at t0 for individual cell simulations (d) and group simulations (h). The EMT-like cell is marked in red. e-g, i, Diagrams depicting the three types of outcome that were monitored during simulations, examples are shown for single cell simulations (e-g) and groups (i) with loss of apical adhesion (***A***). INM, interkinetic movements; M, mitosis; PRAM, pre-mitotic rapid apical movements.

In this study, we define four computational EMT-like events named ***A***, ***B***, ***S*** and ***P*** as follows (Figure 1c). ***A*** is the loss of apical adhesion. When ***A*** occurs, a cell detaches its apical point from its neighbors. The neighbors heal the local “wound” by creating a new apical-apical bound. ***B*** is the detachment of the basal point from the basal line. When ***B*** occurs, the basal point can be displaced within the whole 2D plane of the model. By default, after detachment, basal points and nucleus-to-basal springs remain passive. A seek-and-grab behavior of basal points can be further implemented and is called ***P*** for protrusion. ***P*** is a secondary event that can only occur if ***B*** happened. If ***P*** occurs, the basal spring actively extends until it reaches the area beneath the basal line. When it does, the basal point attaches and pulls, generating a traction force on the nucleus via the nucleus-to-basal spring. The relaxation of the straightness spring is called ***S*** and no longer imposes that apical and basal points and the nucleus be positioned along a straight line. Biologically, this can be interpreted mostly in two ways: i) a mild relaxation of apicobasal polarity, ii) as a loss of stiffness. Indeed, after relaxation of the straightness spring, a cell offers less resistance to lateral pressure (e.g. local crowding, moving nuclei due to INM). Finally, INM can be turned ON or OFF in EMT cells or in control cells.

To assess the impact of simulated scenarios, we monitored three main outputs: i) apical and ii) basal extrusions, which respectively correspond to the nucleus of a cell being located beyond the apical domain or the basal line of the epithelium, and iii) basal positioning (Figure 1d-i). In this last situation, a cell still has its nucleus within the confines of the epithelium but it is located basally with respect to the mean position of the other nuclei in the tissue. This is a relevant factor since it has been shown that, in the case of chick neural crest cell delamination, 90% of the delaminating cells had their nuclei located basally within the hour preceding extrusion^20^. A perfect physiological situation (e.g. neural crest delamination) would be 100% of basal extrusion and 0% of apical extrusion. Apical extrusion is considered a failure, as cells extruding on the apical side would die due to lack of survival signals and would not be able to access extracellular matrix to migrate.

The time scale of the model was previously calibrated using cell cycle length and the normal growth of the chick neuroepithelium^29^. Hereafter, the timings mentioned for simulations correspond to the equivalent biological time. All simulations run for 6 hours to allow the epithelium to reach a steady state before EMT-like events start being implemented. To avoid all simulated events to occur simultaneously and allow for various order of events, there is a 6 to 24h window of opportunity for EMT events to occur which is represented as a faint yellow area on each graph with a given output as a function of time. The simulations then run for an extra 30h (54 hours of total simulation time) so that the consequences of the events can be monitored. This schedule is in agreement with the time frame of chicken head and trunk neural crest cell delamination^10^. By default, and unless otherwise stated, all control cells in the tissue undergo INM while INM in EMT cells is on or off as indicated on the figures. All settings of sEMtor corresponding to each simulation of the study are in S1_Table.

### One-event EMT-like scenarios in individual cells or groups, with or without INM

To assess the impact of EMT-like events, we started by simulating one event (***A***, ***B***, or ***S***) in one EMT cell with INM on. ***A*** leads to apical extrusion of some cells during the permissive EMT period (Figure 2a, grey curve) and this proportion no longer increases after the end of the EMT time window (Movie S1, Figure 2a, grey curve plateaus after 24h). This suggests that cells that extrude apically are in a specific situation at the time of ***A***. Indeed, these cells have their nuclei located apically when ***A*** occurs (Figure S1). ***A*** also leads to a progressive increase of basal positioning (Figure 2b, grey curve). Very few of these cells eventually extrude basally (Figure 2c, grey curve). By contrast, ***B*** has no immediate effect but progressively leads to apical extrusion (Figure 2a, black curve). Finally, ***S*** is not sufficient to promote extrusion on either side or basal positioning of nuclei in the tissue (Figure 2a-c, dotted line, overlapping with the X axis). Simulating the same scenarios in a cluster of 11 cells (Figure 2d-f) leads to the same outcomes with a slight increase in the apical (Figure 2d) and basal (Figure 2f) extrusion rates (Movie S1 and S2).

**Figure 2.**
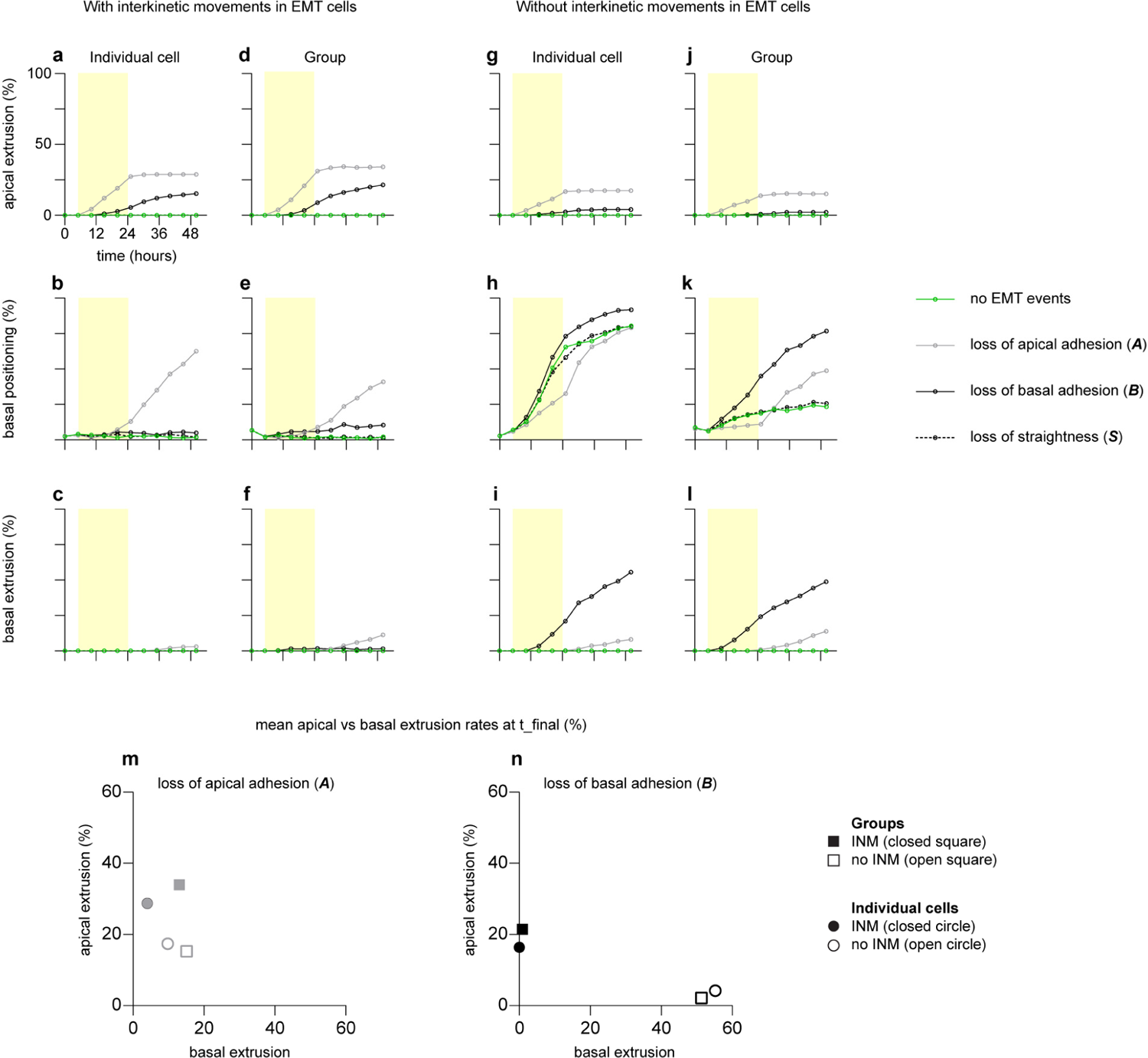
Rates of apical and basal extrusion in one-event EMT-like scenarios a-l, Rates of apical/basal extrusion and basal positioning after simulating one EMT-like event in cells with interkinetic movements (a-f) either individually (a-c) or in groups (d-f) or in cells without interkinetic movements (g-l) either individually (g-i) or in groups (j-l). The EMT-like events are loss of apical adhesion (***A***, grey lines), loss of basal adhesion (***B***, black lines), loss of straightness (***S***, dotted lines). Data for control cells with no EMT-like event are plotted in green. Yellow area on each graph represents the time window of opportunity for EMT-like events to occur (see main text). Each simulation n = 500 for individual cells, n = 50 for groups of 11 cells. All graphs from panels a to l have the same X and Y axes scales, as labelled in a. Note that ***A*** (b, e) or loss of INM (h, k) lead to basal positioning while a combination of ***B*** and loss of INM leads to basal extrusion (i-l). m-n, Scatter plots of rates of apical (Y axis) and basal (X axis) extrusion at t_final for scenarios with ***A*** (m) or ***B*** (n). Note that loss of INM (going from closed to open symbols) has more impact than passing from individual cells to groups (going from circles to squares). Also note that loss of INM has more impact when combined with ***B*** than with ***A***.

Next, to examine the role of INM, we repeated the same simulations without INM in the EMT cells (Figure 2g-l, Movie S3). In individual cells, cancelling INM has dramatic effects. It reduces apical extrusion rates (Figure 2g) and is sufficient to favor basal positioning, even when no other event is taking place (Figure 2h, green curve). Interestingly, the loss of INM synergizes with the other events to increase basal extrusion, mildly in association with ***A*** (Figure 2i, grey curve), dramatically in association ***B*** (Figure 2i, black curve). The same simulations in groups (Figure 2j-l) show the same trends for extrusion. Thus, loss of INM promotes basal positioning and basal extrusion.

Since ***A*** or ***B*** lead to extrusion on either side, to better compare the relative efficiency of each scenario we plotted the apical and basal extrusion rates at t_final as scatter plots (Figure 2m-n). In the case of ***A*** with INM (Figure 2m) going from a single cell (closed circle) to a group (closed square) increases both apical and basal extrusion rates. By contrast, in absence of INM going from a single cell (open circle) to a group (open square) slightly improves the efficiency with less apical and more basal extrusion. In the case of ***B*** (Figure 2n), the main effect is seen when turning off INM. While no basal extrusion is observed in cells performing ***B*** with INM (closed circle and square), turning off INM (open circle and square) acts as a switch that reduces apical extrusion and dramatically enhances basal extrusion when ***B*** occurs.

INM prevents basal positioning of nuclei cell autonomously by bringing nuclei apical at each round of G2/M. However, it also favors apical crowding^29^ and thus might act non-cell autonomously to promote basal positioning of nuclei outside of the G2/M phases. If this is correct, canceling INM in normal cells should reduce the rates of basal positioning and basal extrusion of EMT cells. Thus, we repeated simulations corresponding to Figure 2g-l but this time without INM in both EMT and control cells (Figure S2). Indeed, the rates of basal positioning and basal extrusion of EMT cells were dramatically reduced after INM was canceled in normal cells. These data indicate that INM in normal cells can influence basal extrusion of EMT cells in a non-cell autonomous manner.

### Two-event EMT-like scenarios in individual cells or groups, with or without INM

Next, we simulated two-event EMT scenarios in individual cells or groups. For simplicity, we plotted only the rate of apical and basal extrusion on Figure 3 (Figure 3a-d). Rates of basal positioning are provided in Figure S3. In individual cells, ***S*** had no major effect when coupled with ***A*** or ***B*** (Figure 3a-b, black and grey curves) compared to ***A*** or ***B*** alone. By contrast, coupling ***A*** and ***B*** lowered apical extrusion and increased the rate of basal extrusion (Figure 3a-b, brown curves) compared to ***A***. Performing the same simulations in groups (Figure 3c-d), increased further the rate of basal extrusion. If the same two-event scenarios are now implemented in cells where INM are cancelled (Figure 3e-h), efficiency is further improved with less apical and more basal extrusion. This is especially true for the scenarios in which ***B*** occurs but not ***A***. Interestingly, the combination of loss of INM and ***B*** (Figure 2i-l, black curves) is more efficient at promoting basal extrusion than coupling ***A*** and ***B*** in any order (Figure 3f-h, brown curves).

**Figure 3.**
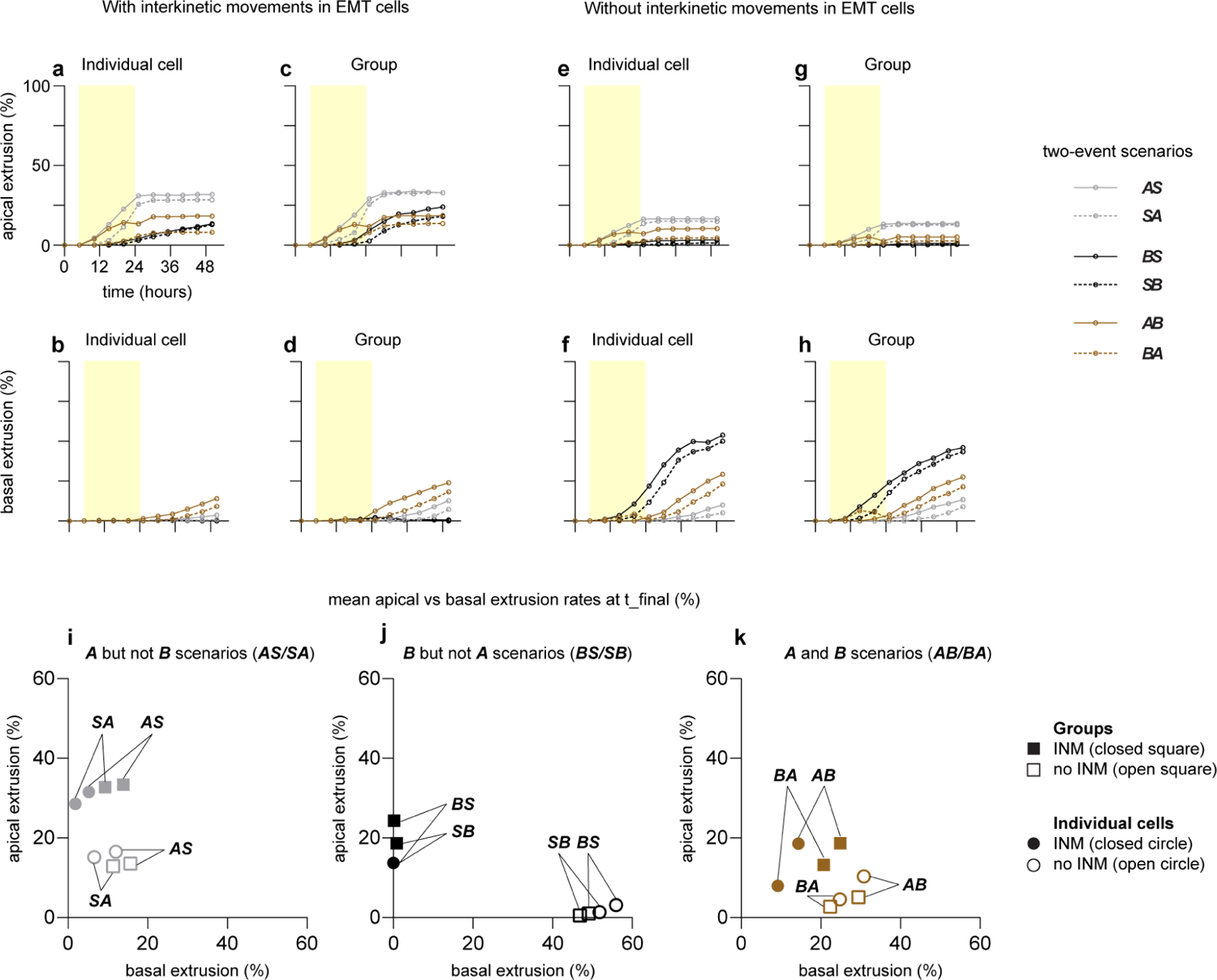
Rates of apical and basal extrusion in two-event EMT-like scenarios a-l, Rates of apical/basal extrusion after simulating two EMT-like events in cells with interkinetic movements (a-d) either individually (a-b) or in groups (c-d) or in cells without interkinetic movements (e-h) either individually (e-f) or in groups (g-h). The EMT-like scenarios are ***AS*** (grey lines), ***SA*** (grey dotted line), ***BS*** (black line), ***SB*** (black dotted line), ***AB*** (brown line), ***BA*** (brown dotted line). Yellow area on each graph represents the time window of opportunity for EMT-like events to occur. Each simulation n = 500 for individual cells, n = 50 for groups of 11 cells. All graphs from panels a to h have the same X and Y axes, as labelled in a. Note that loss of INM reduces apical extrusion and increases basal extrusion for all scenarios. i-k, Scatter plots of rates of apical (Y axis) and basal (X axis) extrusion at t_final for scenarios with ***A*** but not ***B*** (***AS/SA***, i), with ***B*** but not ***A*** (***BS/SB***, j), with ***A*** and ***B*** (***AB/BA***, k). Note that, in absence of INM (open symbols), the occurrence of both ***A*** and ***B*** does not have a cumulative effect on the rate of basal extrusion. Without INM (open symbols), scenarios with ***A*** and ***B*** lead to more extrusion than with ***A*** alone but less than with ***B*** alone.

Plotting the final extrusion rates as scatter plots (Figure 3i-k), better shows the various trends. In particular, we note that ***AB*** scenarios are less sensitive to loss of INM and the size of the EMT population (Figure 3k) than scenarios in which ***A*** or ***B*** occur separately (Figure 3i-j). These plots also help to appreciate the modulating effect of ***S*** as in scenarios where ***S*** happens after ***A*** showing a slight increase of basal extrusion. Finally, these data also indicate that ***A*** partially cancels the effect of losing INM. Indeed, loss of INM in the ***AB*** scenarios has a less dramatic effect than in scenarios with only ***B***. In a cell with no attachment to the apical surface, INM can no longer bring the nucleus apically and thus performing INM or not becomes less relevant.

### Three-event EMT-like scenarios in individual cells or groups, with or without INM

Following the same logic, we ran three-event scenarios with ***A***, ***B*** and ***S*** in various orders: with INM, in individual cells (Figure 4a-b) or groups (Figure 4c-d); without INM in individual cells (Figure 4e-f) or groups (Figure 4g-h). Rates of apical and basal extrusions are on Figure 4 and rates of basal positioning are shown in Figure S3. Interestingly, the scenarios in which ***A*** occurs before ***B*** and is the first event (Figure 4a-h, brown curves) are systematically above all other scenarios in all tested conditions. It should be noted here that it is true for both apical and basal extrusions indicating that this order of event favors extrusion in general and not basal extrusion in particular.

**Figure 4.**
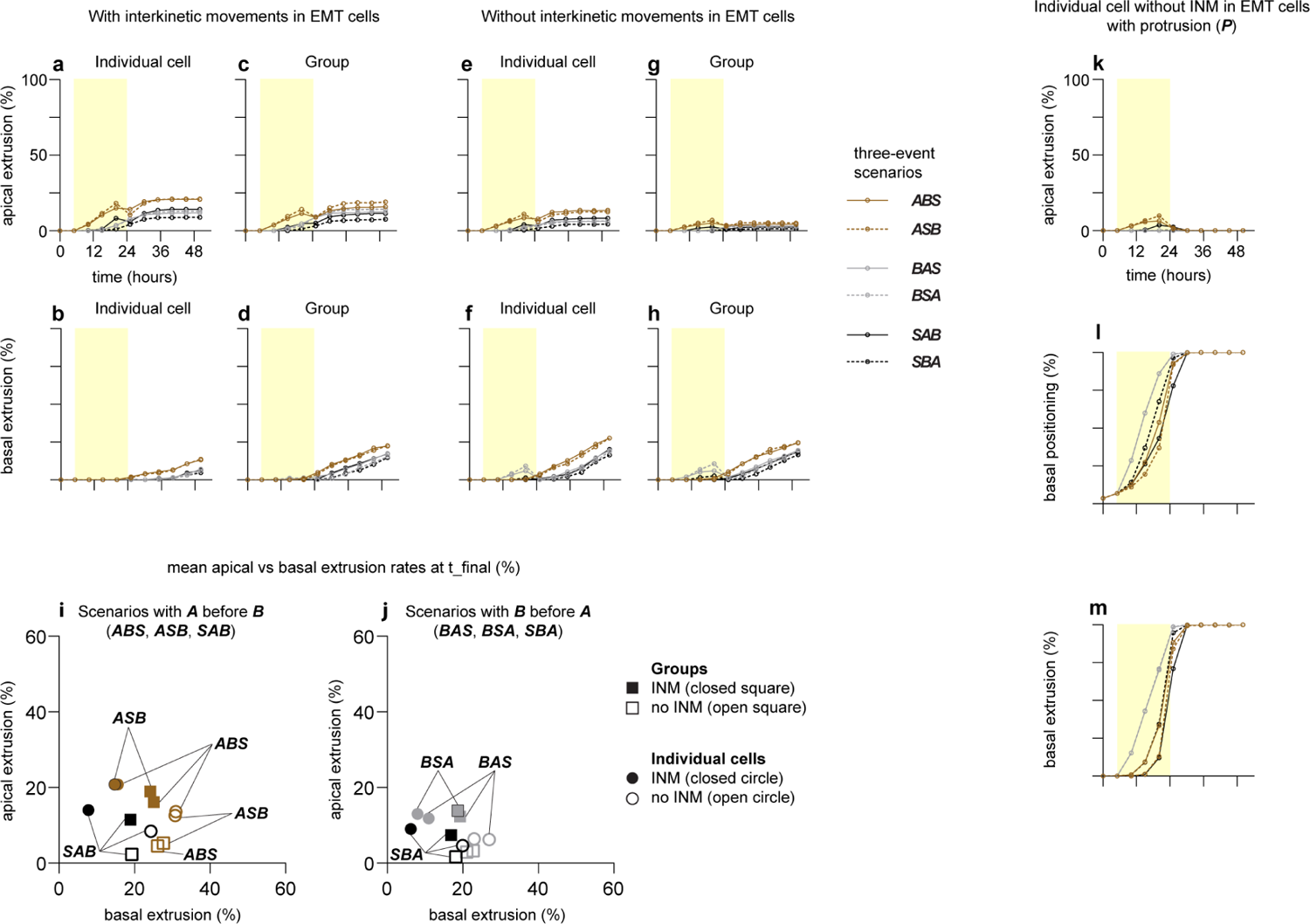
Rates of apical and basal extrusion in three-event EMT-like scenarios a-j, Rates of apical/basal extrusion after simulating three EMT-like events in cells with interkinetic movements (a-d) either individually (a-b) or in groups (c-d) or in cells without interkinetic movements (e-h) either individually (e-f) or in groups (g-h). The EMT-like scenarios are starting with ***A*** (***ABS***, brown lines; ***ASB***, brown dotted lines), starting with ***B*** (***BAS***, grey lines; ***BSA***, grey dotted line) or starting with ***S*** (***SAB***, black lines; ***SBA***, black dotted lines). Yellow area on each graph represents the time window of opportunity for EMT-like events to occur. Each simulation n = 500 for individual cells, n = 50 for groups of 11 cells. All graphs from panels a to h have the same X and Y axes, as labelled in a. Note that loss of INM reduces apical extrusion and increases basal extrusion for all scenarios. i-j, Scatter plots of rates of apical (Y axis) and basal (X axis) extrusion at t_final for scenarios with ***A*** before ***B*** (***ABS, ASB, SAB***, i) or with ***B*** before ***A*** (***BAS***, ***BSA***, ***SBA***, j). Note that the order of events seem to only have moderate effect on the rates of apical or basal extrusion. k-m, Rates of apical extrusion (k), basal positioning (l) and basal extrusion (m) of individual EMT cells in three-event EMT-like scenarios without INM ***with*** implementation of protrusive-like behavior (***P***); from 500 simulations. Note the reduction of apical extrusion in panel k compared to panel e, and the increase of basal extrusion in panel m compared to panel f. Rates of basal positioning in panel l should be compared to those in Figure S3, panel g.

Plotting the final extrusion rates as scatter plots (Figure 4i-j), helps to better appreciate the variations induced by the order of events, population size (single vs group) and occurrence of INM. One striking observation is that scenarios with all events (***ABS*** in any order and no INM), which might represent a complete EMT, are comparatively less efficient at producing basal extrusion than some associations such as ***B*** and no INM that might be considered as partial EMT scenarios from a biological stand point.

Collectively, these simulations with one, two or three events with or without INM show that: 1/ more than one scenario can lead to basal extrusion, 2/ there is no major group effect as the outcome of a given scenario can be seen in individual cells and does not dramatically change if simulated in a group of cells, 3/ the position of the nucleus at the time of epithelial destabilization is a major factor of the directionality of extrusion, 4/ when occurring in absence of INM, scenarios with ***B*** but not ***A*** are more efficient than scenarios with ***A*** and ***B***, but when both events do occur the rate of extrusion is higher when ***A*** occurs before ***B***.

Importantly, we observed that none of the above scenarios recapitulates the biological situation observed during physiological EMT. ***In vivo***, neural crest cells either leave the neuroepithelium via basal extrusion or remain in the dorsal neural tube. There is virtually no apical extrusion. The same observation is true for gastrulating mesoderm. By contrast, in the model, all scenarios tested so far lead to some degree of apical extrusion. This suggests that, biologically, something prevents apical extrusions from occurring and/or that something strongly biases extrusion towards the basal side such that apical extrusions of neural crest cells are rare. A second striking difference is the timing of extrusion. In the best case scenario (***B*** without INM in an individual cell), the rate of basal extrusion after 54h of simulated biological time only reaches 50% (Figure 2i, grey curve). ***In vivo***time-lapse imaging shows that neural crest cells usually take no more than a few hours to leave the neuroepithelium^20, 21^.

The basally oriented force generated by apical crowding of nuclei due to INM in normal cells is sufficient to displace nuclei of EMT cells basally if EMT cells have lost their own INM or performed ***A*** (as shown in Figure 2b, 2h and Figure S1). However, this is not enough to ensure a timely exit of cells. The dichotomy between modelling and biological data strongly suggests that an actual driving force is needed. The most obvious candidate for the task is a protrusive activity directed towards the basal compartment. It is known that neural crest cells upregulate multiple integrin subunits implicated in cell motility prior to extrusion^30, 31^. In the model, protrusive activity is represented by ***P***, a seek-and-grab behavior of the basal point that can only occur if ***B*** previously took place. We rerun the EMT scenarios with three events in an individual cell without INM and added ***P*** (Figure 4k-m, Movie S4). Adding ***P*** dramatically enhances the rate of basal extrusion and shortens the time between the start of EMT-like events and extrusion. Importantly, it also suppresses apical extrusion. Both trends are observed in group simulations as well (Movie S4). Collectively these data indicate that a timely basal exit requires a destabilization of the epithelial structure (***B***, or ***AB***) coupled with a basal positioning of the nuclei (i.e loss of INM) and an actual driving force towards the basal compartment (***P***).

### Simulation of heterogeneous clusters of EMT cells

Next, we wondered if implementing heterogeneity, with neighboring cells performing different EMT-like scenarios, would affect the relative efficiency of the various scenarios. We performed simulations with groups of cells but, this time, we implemented a gambling routine at the onset of simulations that attributes random times of occurrence for ***A***, ***B*** and ***S*** for each cell of the group (see Supplementary Information). For all permutations of one, two and three-event scenarios to be statistically possible, we set the probability of picking a time for each event to 70% and that of not picking a time to 30%. Random times are within the time window of opportunity of 6 to 24h. Times for each event are set at the onset of simulation and, if cell division occurs prior to an event, daughter cells inherit the times of the mother cell.

Note that in absence of INM heterogeneity has little to no effect. e-f, Rates of apical (e) and basal (f) extrusion per EMT-like scenarios under various heterogeneous conditions: all EMT-like cells with INM (black triangles), none of the EMT-like cells with INM (open black triangles), 50% of EMT-like cells with INM, 50% without INM (open downward brown triangles), EMT-like cells with 50% chance of having INM and 50% chance of making protrusions (grey diamond). Note that ***P*** dramatically increases the rate of basal extrusion while reducing apical extrusion. g, Scatter plot of the mean rates of apical and basal extrusion across all scenarios for the four heterogeneous conditions presented in panels e and f.

We simulate a group of EMT cells surrounded by control cells on each side. To make sure that each scenario is generated multiple times, we ran fifty thousand simulations. This was done with or without INM in EMT cells. We then plotted apical and basal extrusion rates of each scenario implemented in heterogeneous clusters and compared them with the efficiencies when done in individual cells or homogeneous groups (Figure 5). With INM (Figure 5a-b), for most scenarios, heterogeneity (black triangles) increases extrusion rates. This indicates that, in the context of EMT cells performing INM, heterogeneity increases epithelial destabilization leading to more cells leaving the tissue on either side but does not create a directional bias. When INM is cancelled in EMT cells, the effect of heterogeneity is weak. There is either no effect or slightly less extrusion depending on the scenario (Figure 5c-d). We also plotted the fold difference between extrusion rates going from single cell to homogeneous groups and from homogeneous to heterogeneous groups per scenario and globally (Figure S4) to better appreciate the impact of critical mass and heterogeneity on extrusions. Some scenarios are heavily impacted by going from individual cells to groups such as ***SA*** while others (e.g. ***BS***, ***SB***, ***BAS***, ***BSA***, ***SAB***, ***SBA***) are sensitive to the heterogeneous context.

**Figure 5.**
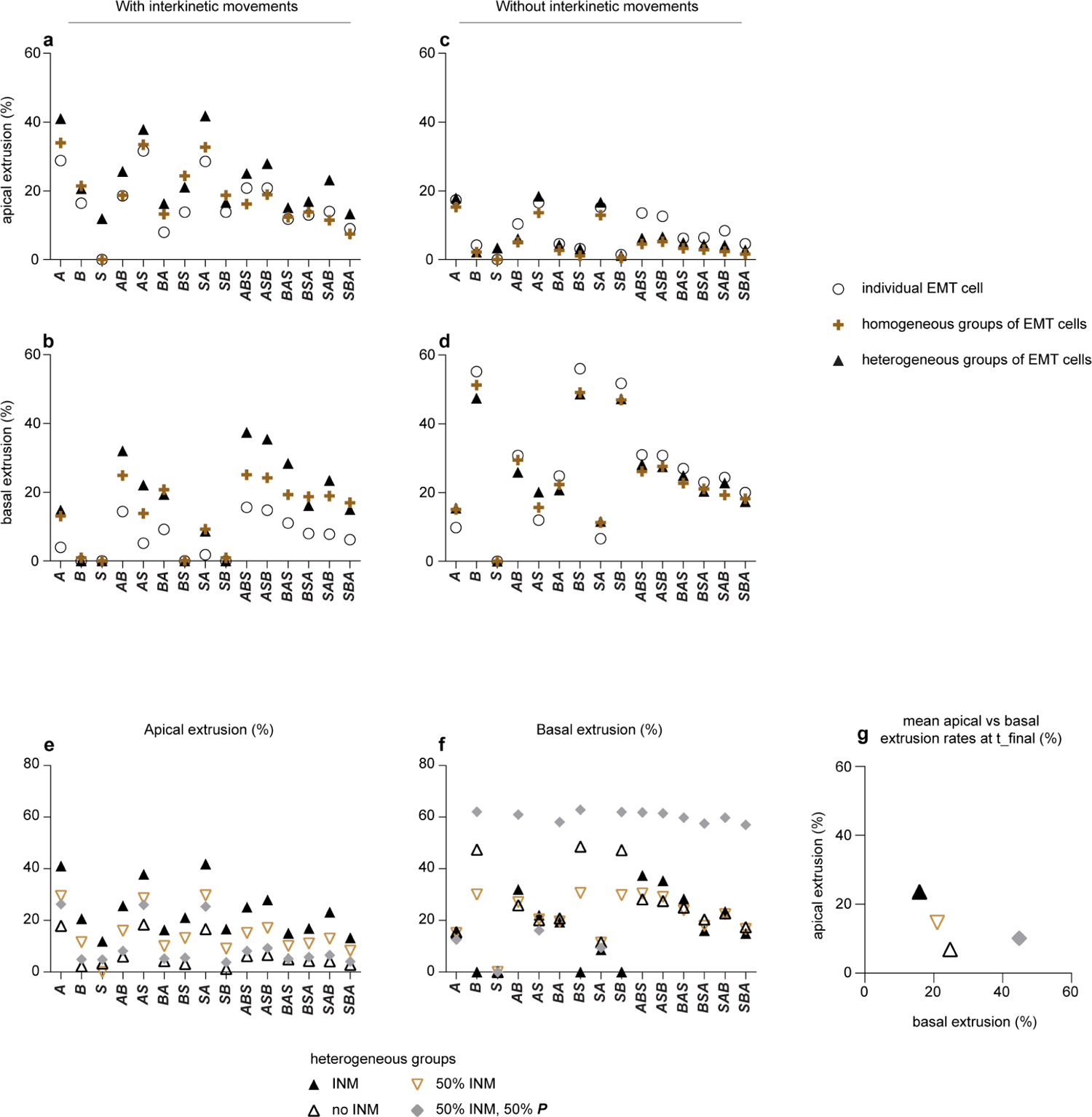
Impact of heterogeneity on the rates of apical and basal extrusion a-b, Rates of apical (a) and basal (b) extrusions per EMT-like scenarios with interkinetic movements implemented in individual cells (open circles), homogeneous groups (brown cross) and heterogeneous groups (black triangle). Note that triangles are above the other symbols for both apical and basal extrusion rates for nearly all scenarios indicating that heterogeneity increases overall extrusion rates. c-d, Rates of apical (c) and basal (d) extrusion per EMT-like scenarios without interkinetic movements implemented in individual cells (open circles), homogeneous groups (brown cross) and heterogeneous groups (black triangle).

To further increase heterogeneity, we allowed cells to choose to perform INM or not during the gambling phase. This generates populations with 50% of the EMT cells with INM and 50% without. We then compared extrusion rates encompassing all scenarios for this simulation with those of the heterogeneous clusters with or without INM in EMT cells (Figure 5e-g). Interestingly, going from all cells with or without INM to a 50-50% situation mostly modulates apical extrusion without affecting the overall efficiency of basal extrusion (Figure 5g). Finally, we introduced ***P*** in the gambling session. To account for the increased number of possible scenarios we ran a hundred thousand simulations. All scenarios with ***P*** are extremely efficient at performing basal extrusion and preventing apical extrusion (Figure 5e-g, grey diamonds).

To further explore the data from the heterogeneous simulations, we ranked all scenarios per efficiency of basal extrusion binning them per time of occurrence of each EMT-like event and per position of their nucleus at the onset of EMT (S1_Spreadsheet). We noticed some interesting trends. As expected, scenarios with ***P*** top the list with a hundred percent efficiency of basal extrusion. This includes partial EMT-like scenarios with one or two events only. More interesting, among the most efficient scenarios without ***P***, we find cells undergoing almost any scenario but sharing the common fact that their nuclei were near the basal side when EMT-like events were initiated. This reinforces the notion that nuclear positioning biases the directionality of extrusion.

In order to assess the relative impact of the different events, we ran correlation analyses between the rate of apical or basal extrusion and the following parameters: i) the occurrence of the events (***A***, ***B***, ***S***, ***P***), ii) the position of the nucleus at the onset of simulation, the onset of EMT or when ***A***, ***B***, ***S*** or ***P*** occur, iii) the timing of events and iv) the time interval between the first EMT event and the last (for scenarios including at least two events). We used the data from the most heterogeneous situation in which EMT cells can perform any scenario with 50% chance of performing INM, and 50% chance of ***P*** (Figure 6). These analyses reveal that ***P*** is the only event whose occurrence is negatively correlated with apical extrusion and positively correlated with basal extrusion. In addition, the position of the nucleus when the first EMT-like event occurs (***y_emt***), when A, B or S occur (***y_A/B/S***) is systematically correlated with extrusion. More precisely, ***y_emt***, ***y_A***, ***y_B*** and ***y_S*** are negatively correlated with basal extrusion meaning that the lowest values of y (basal positions of nuclei) favor basal extrusion while higher values of y (apical nuclei) favor apical extrusion. This is true even for events whose overall occurrence is not correlated (e.g. ***A***, ***S***) with either extrusion. This means that performing ***A*** in itself does not strongly favor basal extrusion but performing ***A*** while the nucleus is basal strongly correlates with basal extrusion. For scenarios with ***B*** and ***P***, the position of the nucleus (***y_P***) no longer correlates with extrusion. The protrusion generates a basally oriented driving force that bypasses the effect of the nucleus position. This global trend is true even if the correlation between ***y_emt*** and extrusion is plotted per scenario (Figure 6c-d). Finally, the timing of the different events (***t_A***, ***t_B***, ***t_S***) and the duration of the EMT-like scenarios (***Δt_emt***) are not correlated with extrusion. In both INM and non-INM cells, the trend is similar with the noticeable exception of the position of the nucleus at the initiation of the simulation (***y_init***) that goes from not being correlated (INM cells) to being negatively correlated with basal extrusion (no INM cells). This is due to the fact that in cells with INM the initial position of the nucleus is not correlated to its position at the time of EMT because INM changes the nucleus position over time. While in cells without INM, the nucleus is statistically more likely to be basal, thus the weak negative correlation between ***y_init*** and basal extrusion in no INM EMT cells.

**Figure 6.**
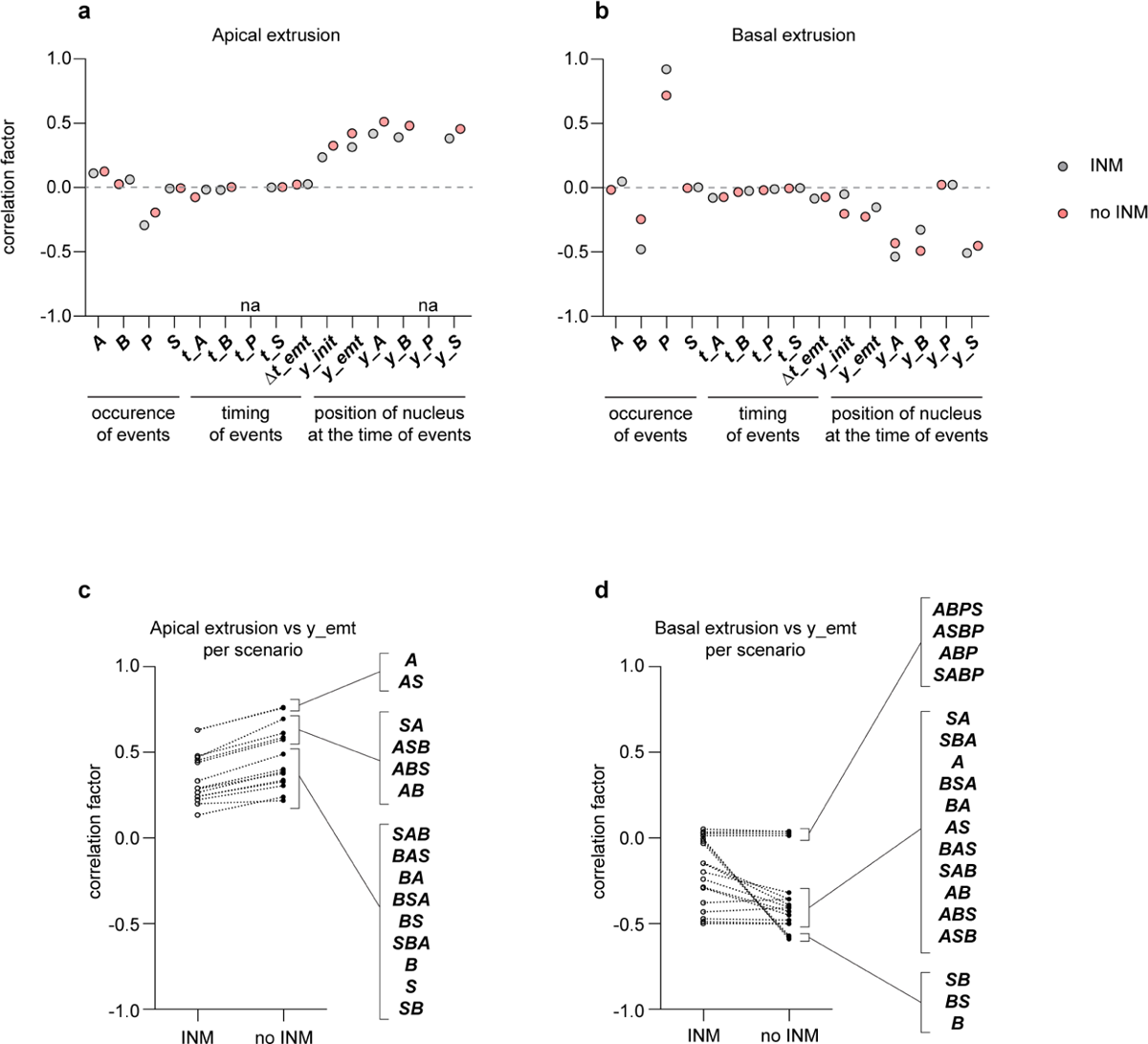
Correlation analyses between the occurrence/timing of events or nuclei position and the rates of apical and basal extrusion in heterogeneous simulations a-b, Correlation factor for a given parameters/event and apical (a) or basal (b) extrusion rates at t_final, either with INM (grey circle), or without INM (pink circles). Correlation factors are plotted for the occurrence of a specific event (***A***, ***B***, ***S*** or ***P***), the timing of a given event (***t_A***, ***t_B***, ***t_S***, ***t_P***), the time interval between the first and last event of any EMT scenarios with more than one event (***Δt_emt***) and the position of a cell’s nucleus at the onset of simulation (***t_init***), the onset of EMT (***y_emt***) or when a given event occurs (***t_A***, ***y_B***, ***y_S***, ***y_P***) across all relevant scenarios. For instance ***B*** occurs in the following scenarios ***B***, ***BS***, ***SB***, ***AB***, ***BA***, ***ABS***, ***ASB***, ***BAS***, ***BSA***, ***SAB*** and ***SBA*** with/without INM and with/without ***P*** but does not occur in ***A***, ***S***, ***AS***, ***SA***. Thus, correlation analyses reflect on the influence of ***B*** across all relevant scenarios. Same logic applies for all parameters tested. Note that the position of the nucleus is positively correlated with apical extrusion and negatively with basal extrusion. Also note that ***P*** is highly positively correlated with basal extrusion and negatively with apical extrusion. c-d, Correlation factor for the position of nuclei when the first emt event occurs (***y_emt***) and apical (c) or basal (d) extrusion per scenarios with or without INM. Note that apical extrusion (c) is systematically positively correlated with nucleus position (apical) regardless of the scenario and INM status. By contrast, note for instance that basal extrusion and nucleus position (d) can show no correlation (scenarios with ***P***) or display strong negative correlation (scenarios with ***B*** but neither ***A*** nor ***P*** without INM).

Overall, the simulations with heterogeneous populations and correlation analyses indicate that i) heterogeneity can act as a destabilization factor increasing extrusion rates on either side of the epithelium and further support the idea that ii) the position of the nucleus and protrusion heavily influence the directionality and timing of extrusion. Next, we decide to confront our ***in silico*** observations to the physiological EMT of neural crest cells or to the destabilization of epithelial features in the neuroepithelium.

### Regulation of INM is looser in the neural crest domain than in the rest of the neural tube

Simulations indicate that the basal positioning of nuclei is crucial for basal extrusion. This is consistent with data from trunk neural crest cells suggesting that synchronizing EMT with the S-phase of the cell cycle represent a window of opportunity for cells to exit the neural tube while their nuclei are basal^32^. It is also in agreement with the observation that 90% of neural crest cells have their nuclei basally positioned in the hour preceding delamination^20^. Our simulations indicate that another way to increase the probability of having a basal nucleus is to cancel INM. Lack of tight regulation of INM leads to non-apical mitoses that can be easily observed by immunostaining on fixed samples. Previous data showing that non-apical mitoses are frequent in the trunk neural crest domain at the time of EMT suggests the absence of a tight regulation of INM during this process^33^. We then wondered whether such non-apical mitoses were a consequence of EMT itself (e.g. apical detachment) or if they could be observed in the neural crest domain prior to EMT. To this end, we monitored the distribution of mitotic cells using phospho-Histone H3 staining in cephalic and trunk neural crest regions prior and during EMT (Figure 7a-b). In pre-EMT neural crest cells at cephalic (Figure 7c-e) and trunk levels (Figure 7f-h) the rate of non-apical mitoses is higher than in the neuroepithelium (where no EMT occurs). Interestingly, the rate of non-apical mitoses in the neuroepithelium significantly drops from posterior to anterior regions (Figure 7g) indicating that a tight regulation of INM is progressively implemented as the neural tube develops. The neural crest domain follows an opposite trend with an increase of non-apical mitoses as EMT is initiated. Overall, these data indicate that a significant amount of non-apical mitoses is observed before the onset of EMT showing that neural crest cells initially lack a tight regulation of INM prior to EMT implementation and that EMT further increases the rate of non-apical mitoses. According to our simulations, this would favor basal positioning of nuclei, which may facilitate basal, rather than apical, extrusion of neural crest cells upon epithelial destabilization.

**Figure 7.**
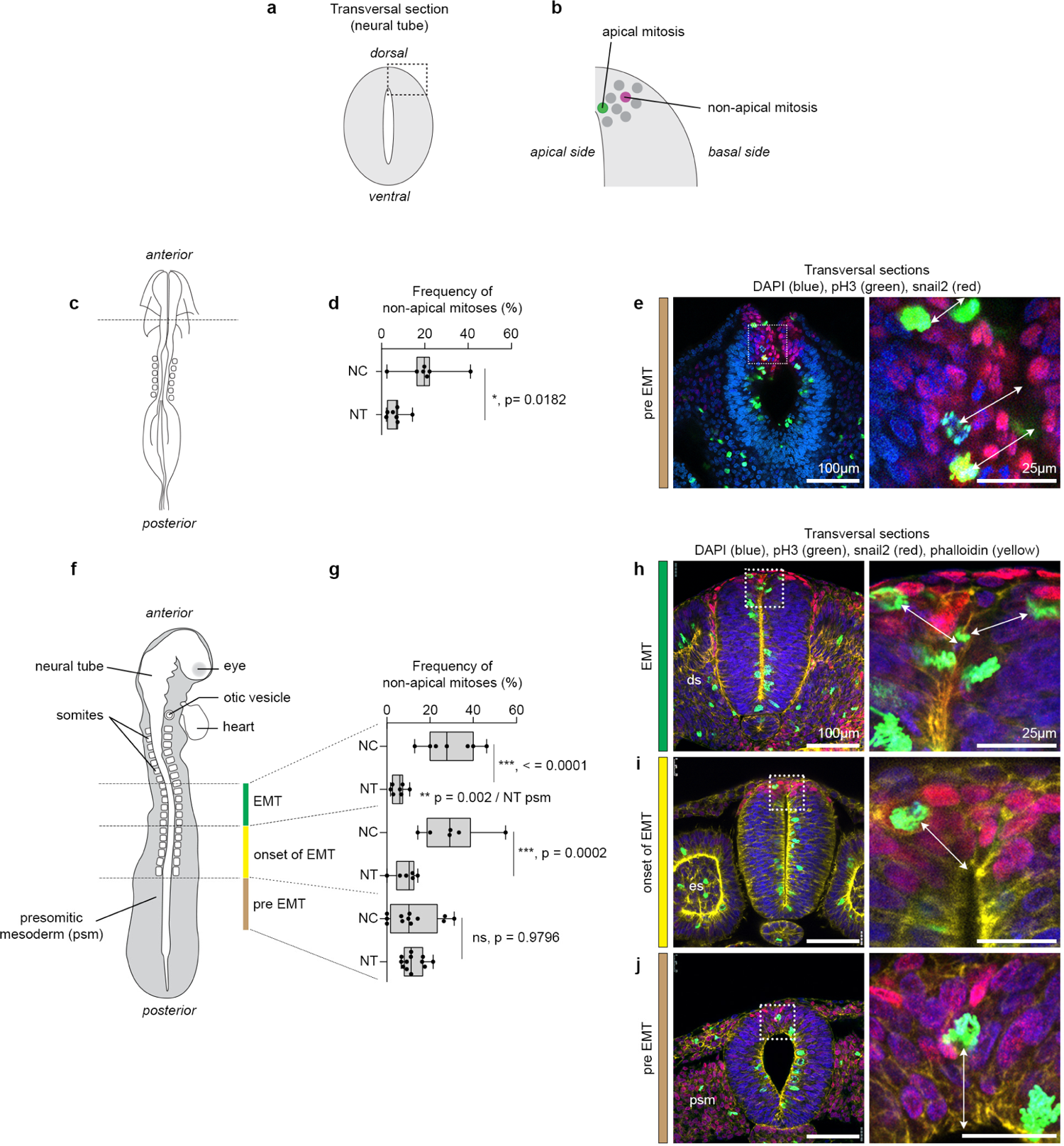
Neural crest territory has a high rate of non-apical mitoses before the onset of EMT a-b, Diagrams representing transversal section of the neural tube and how we classified apical vs non-apical mitoses. A mitosis is considered apical (green) if there are no other nuclei in between that mitotic figure and the apical domain. It is considered non-apical (magenta) if at least one nucleus separates the mitotic nucleus from the apical side. c-h, Quantification of non-apical mitoses, as percentage per embryo, in neural crest cells and the adjacent neural tube at the level of pre-EMT cephalic neural crest cells (c-e) and at the levels of pre-EMT (neural tube facing presomitic mesoderm (psm), j), early EMT (i) and late EMT (h) trunk neural crest cells. The statistical tests used are: unpaired t-test with Welch’s correction (d, n = 7 embryos per condition), one-way ANOVA/Fisher’s LSD test (g, nembryos = 12 (pre-EMT/psm level), 6 (onset of EMT/-1 to-6 somites), 7 (EMT/-7to-12 somites)).

### Upregulation of integrins contributes to epithelial destabilization

Our simulations point to a critical role of protrusions to ensure a timely and directional extrusion. Interestingly, α4 and α5 integrins are specifically upregulated in neural crest cells prior to delamination^30, 31^. Their inhibition in neural crest cells leads to migration defects^30, 31, 34, 35^ indicating that these integrin subunits are critical for neural crest motility. In addition, some neural crest cells end up located in the lumen of the neural tube indicating apical extrusion. This shows that impairing interaction with the matrix leads neural crest cells undergoing EMT to randomly exit apically or basally. This reinforces the notion that protrusive activity contributes to the directionality of extrusion and not just migration post extrusion. Another striking observation from these studies is the timing of expression of α4 and α5 integrins which start being expressed several hours before any membrane extensions are described. Given that cell-cell and cell-matrix adhesion complexes tend to be spatially segregated and functionally antagonistic^25, 36, 37^, we wondered whether the early upregulation of α4 and α5 integrins could contribute to epithelial destabilization in addition to their role in motility. To test that, we generated expression vectors for chicken α4 and α5 integrins and overexpressed them in the neuroepithelium (Figure 8). 24 hours post electroporation, we analyzed the distribution of multiple apical and basal markers: atypical Protein Kinase C (aPKC), N-cadherin, Pericentriolar Material 1 (PCM1), Laminin and Fibronectin. In embryos overexpressing a control membrane-bound GFP, no defects are observed for any of the markers (Figure 8a, 8e, Figure S5). By contrast, expressing either α4 or α5 or a combination of both was sufficient to lead to ectopic localization of apical and basal markers as well as driving cell extrusion into the lumen (Figure 8b-k, Figure S5). The overall morphology of the neural tube appears normal and no major rearrangements of cell such as rosettes are observed by contrast to what happens after expression of polarity protein Par3^22, 33^ or pro-EMT factors such as Ets1^12^. These data indicate that, in addition to their role in motility, expression of specific integrin subunits can contribute to epithelial destabilization as part of EMT. However, if their expressions are not coupled to other events promoting acquisition of front-rear polarity to ensure protrusive activity toward the basal compartment such destabilization may lead to apical extrusion as we observed.

**Figure 8.**
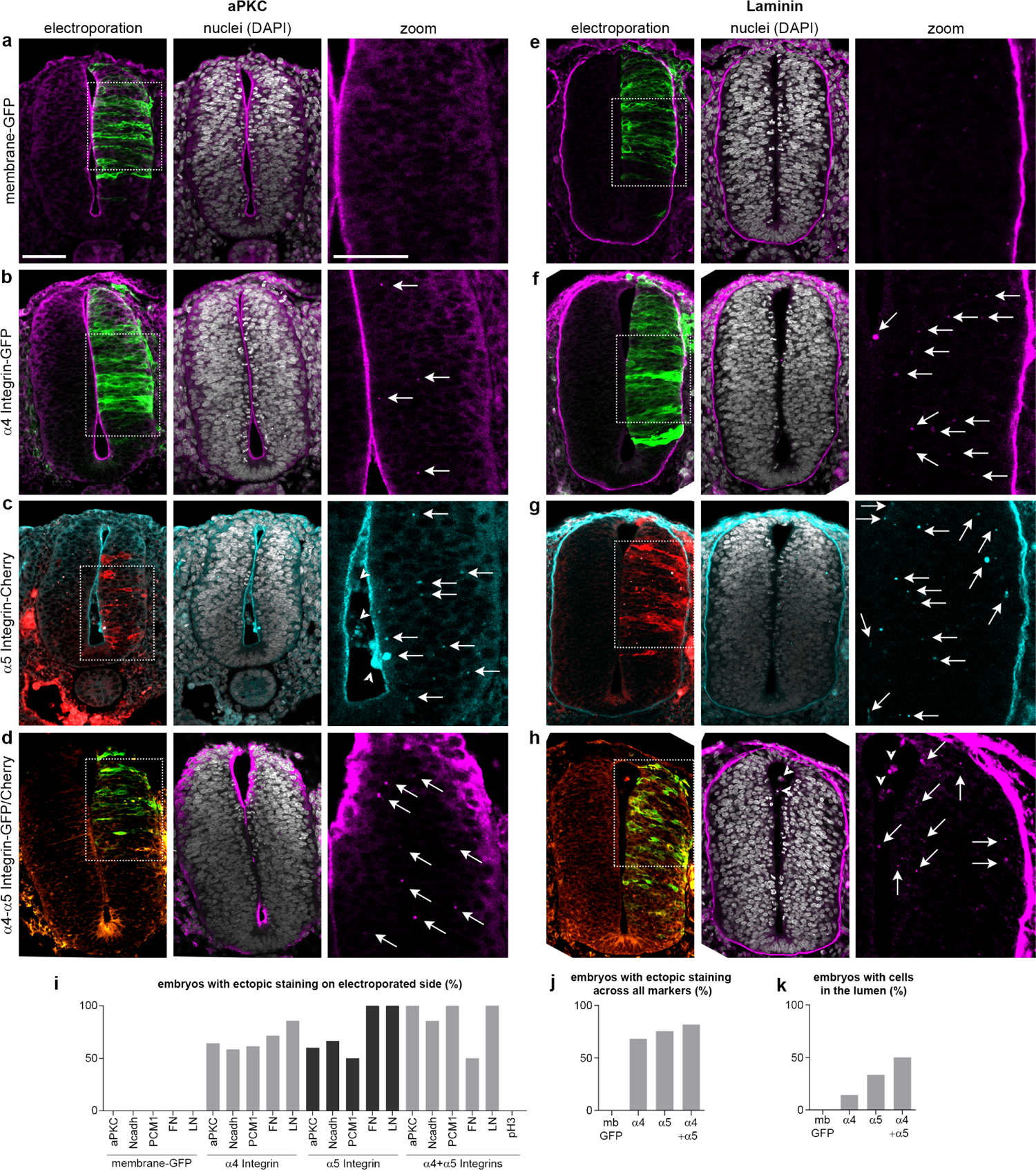
Overexpressions of α4 and α5 Integrins lead to moderate apicobasal defects and apical extrusions a-d, Representative images for immunostaining on cryosections against atypical Protein Kinase C (aPKC) from embryos expressing membrane-GFP (a, n = 3), α4-Integrin (b, n = 14), α5-Integrin (c, n = 5) or a combination of α4 and α5 Integrins (d, n = 7). e-h, Representative images for immunostaining on cryosections against Laminin from embryos expressing membrane-GFP (e, n = 3), α4-Integrin-GFP (f, n = 14), α5-Integrin-Cherry (g, n = 6) or a combination of both α4 and α5 Integrins (h, n = 7). Nuclei are stained with DAPI (grey), α4 and α5 are displayed in green and red respectively. Immunostainings are shown in magenta (a-b, d, e-f, h) or un cyan (c, g). Arrows indicate examples of ectopic staining in the electroporated area. Arrowheads show example of cells that performed apical extrusion. Scale bars in low magnification 80 µm, in zooms 50 µm. i-j, Percentages of embryos with ectopic staining on the electroporated side per marker per experimental condition (i), across all markers per experimental condition (j). k, Percentages of embryos with apical extrusion (cells located in the lumen). Embryos: α4 (n = 14), α5 (n = 9), α4+α5 (n = 8). Representative images of embryos for Fibronectin, Pericentriolar Material (PCM) 1 and N-cadherin are shown on Figure S5.

Along these lines, our simulations also point to the importance of detaching cells from the basal line, as simulations with ***B*** and no INM have a high rate of basal extrusion. Gaps in the basement membrane are often proposed as opportunities for cells to exit the epithelium but their ability to specifically promote basal extrusion is unclear. We tested this in embryos by treating whole trunk explants with Dispase II to partially degrade fibronectin (see Methods). This leads to a reorganization of the matrix with gaps in the laminin surrounding the neural tube (Figure S6). Interestingly, the neuroepithelium is disorganized with local buckling as previously observed with similar treatments at later stages or when affecting actomyosin^29, 38^. It also leads to non-apical mitoses and to protrusive activity on both sides of the epithelium, as well as local loss of N-cadherin and apical extrusion (Figure S6). These data indicate that impairing interaction with the matrix contributes to epithelial destabilization and promotes extrusion but, similarly to upregulation of integrins, is not sufficient to provide a basal bias.

### Cephalic neural crest cells are more heterogeneous than trunk NC cells

Finally, our simulations indicate that heterogeneity (the co-existence of multiple scenarios) favors destabilization and increases overall extrusion rates. Thus, one might expect regions of massive neural crest departure (mesencephalon) to be more heterogeneous than regions of progressive neural crest departure (trunk). Heterogeneity of neural crest populations has been observed at cephalic and trunk levels by single cell transcriptomics^39-42^ but this technique does not allow to assess spatial heterogeneity. Other techniques such as multiplex RNA detection on sections^43^ showed heterogeneity in terms of gene expression but given that the relationship between RNA and protein levels is poor^44^ it is difficult to predict whether detected heterogeneity at RNA levels reflects actual heterogeneity at protein level.

To compare heterogeneity of cephalic and trunk neural crest cells in terms of EMT effectors at protein level, we performed immunostainings against TFAP2α, Snail2 and Sox9, two by two (Figure 9a-d). Snail2 and Sox9 have been implicated in neural crest EMT^17, 18, 45^ and TFAP2α is known to be upstream of both factors^46^. In addition, all three transcription factors are deemed part of the core neural crest regulatory cluster in chicken neural crest cells^43^. Interestingly, Snail2 and Sox9 only lead to partial epithelial destabilization when overexpressed alone but promote basal extrusion together^47^. Thus differential levels of these two proteins may indicate a diversity of phenotypes along the EMT spectrum. In cephalic regions, neural crest cells strongly expressing a couple the above-mentioned markers (Figure 9a-c, arrows) are scattered across the whole neural crest domain and intermingled with cells expressing mostly one or the other proteins. By contrast, in the trunk, neural crest cells strongly expressing either couple of markers (Figure 9a-c, arrows) are concentrated in the most basal part of the neural crest domain. This indicates a higher degree of spatial heterogeneity in cephalic than in trunk neural crest cells. Further, scatter plots of normalized intensities for each pair of markers in head and trunk neural crest cells (Figure 9e-f) show that cephalic neural crest cells display a wider range of staining intensities of all three markers. Indeed, scatter plots for trunk neural crest cells show that most cells have high expression levels for both proteins (top right corner). Intensities per marker are significantly different in trunk and cephalic neural crest cells (Figure 9g) and coefficient of variation (CV) are systematically higher in cephalic than trunk neural crest cells (Figure 9g). Altogether, these data indicate that the cephalic neural crest population is likely to be more heterogeneous than the trunk population. Thus, heterogeneity level correlates with delamination intensity in the neural crest domain, supporting the role of heterogeneity as a modulator of EMT.

**Figure 9.**
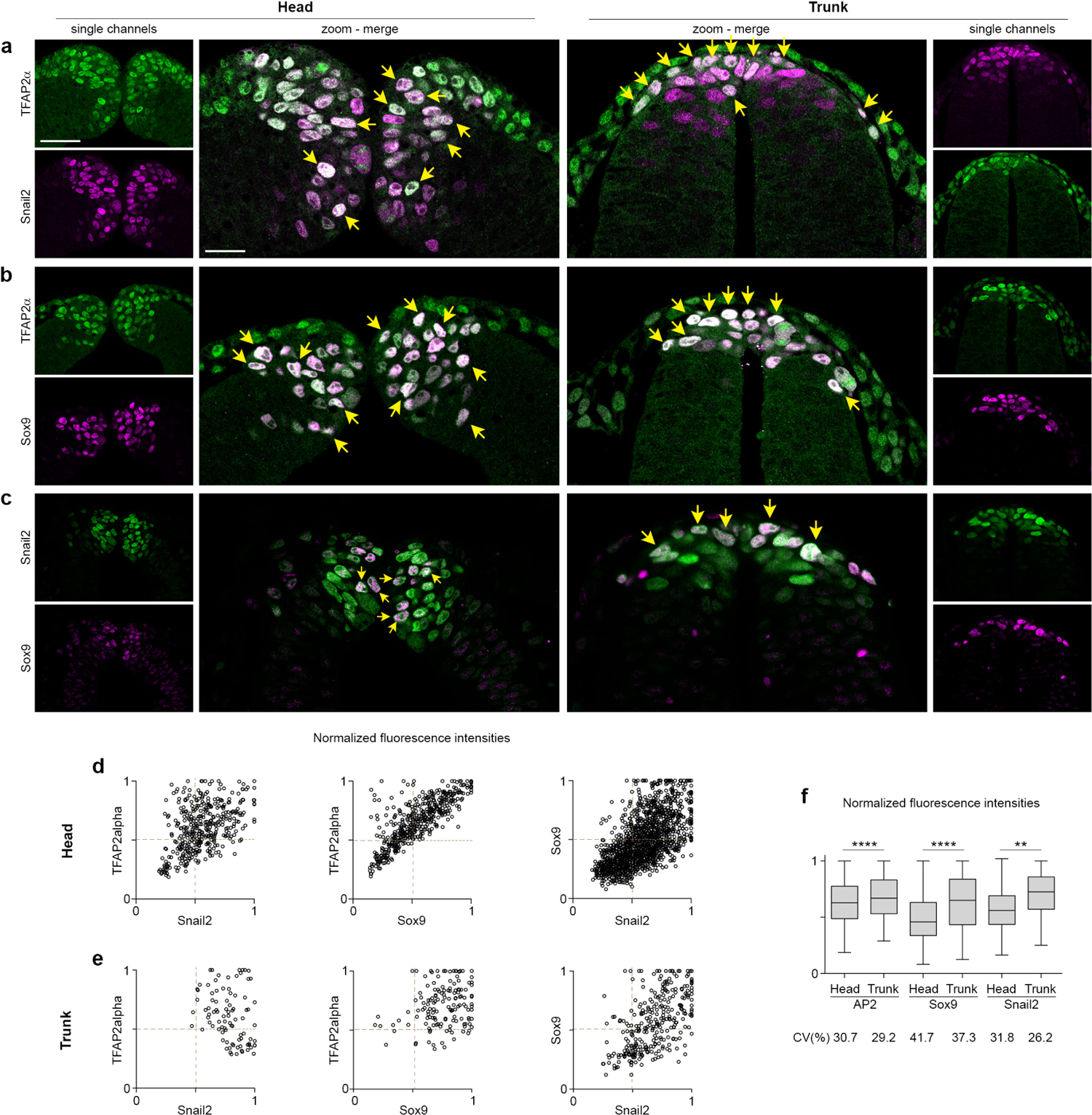
Differential heterogeneity of cephalic and trunk neural crest cells a-c, Double immunostainings against TFAP2α and Snail2 (a), TFAP2α and Sox9 (b), Snail2 and Sox9 (c) on cryosections of cephalic neural crest cells (left columns) or trunk neural crest cells (right column). Scale bars, 50 µm on low magnifications and 25 µm on zooms. Yellow arrows indicate neural crest cells with high staining for both markers. Note that in cephalic regions double-positive cells are scattered while they are basally located in the trunk region. e-f, Scatter plots from double immunostainings, each channel was normalized to its maximum value. g, Box and whisker plot of normalized fluorescence intensity values for each marker at head and trunk levels. Coefficient of variation (CV) are indicated underneath each box plot. Uncorrected Fisher’s LSD, **** p < 0.0001, ** p = 0.011.

In conclusion, rather than a linear cascade of events leading to extrusion followed by cell migration (Figure 10a), our simulated and biological data depict the cellular implementation of EMT as the result of an array of multiple inputs that can cooperate through various scenarios (Figure 10b). This description better fits models of gene regulatory networks of EMT displaying multiple parallel pathways interconnected by feedback loops^48-50^. Importantly, key determinants of timely and directional extrusion appear to be the pre-positioning of the nucleus and protrusive activity. Given that nucleus position is influenced by multiple inputs, heterogeneity can be seen as an emerging property of the system which may further boost epithelial destabilization. Finally, the importance of protrusive activity as an early event to bias the directionality of extrusion strongly suggests that all molecular effectors of the motility machinery may be early rather than late markers of EMT.

**Figure 10.**
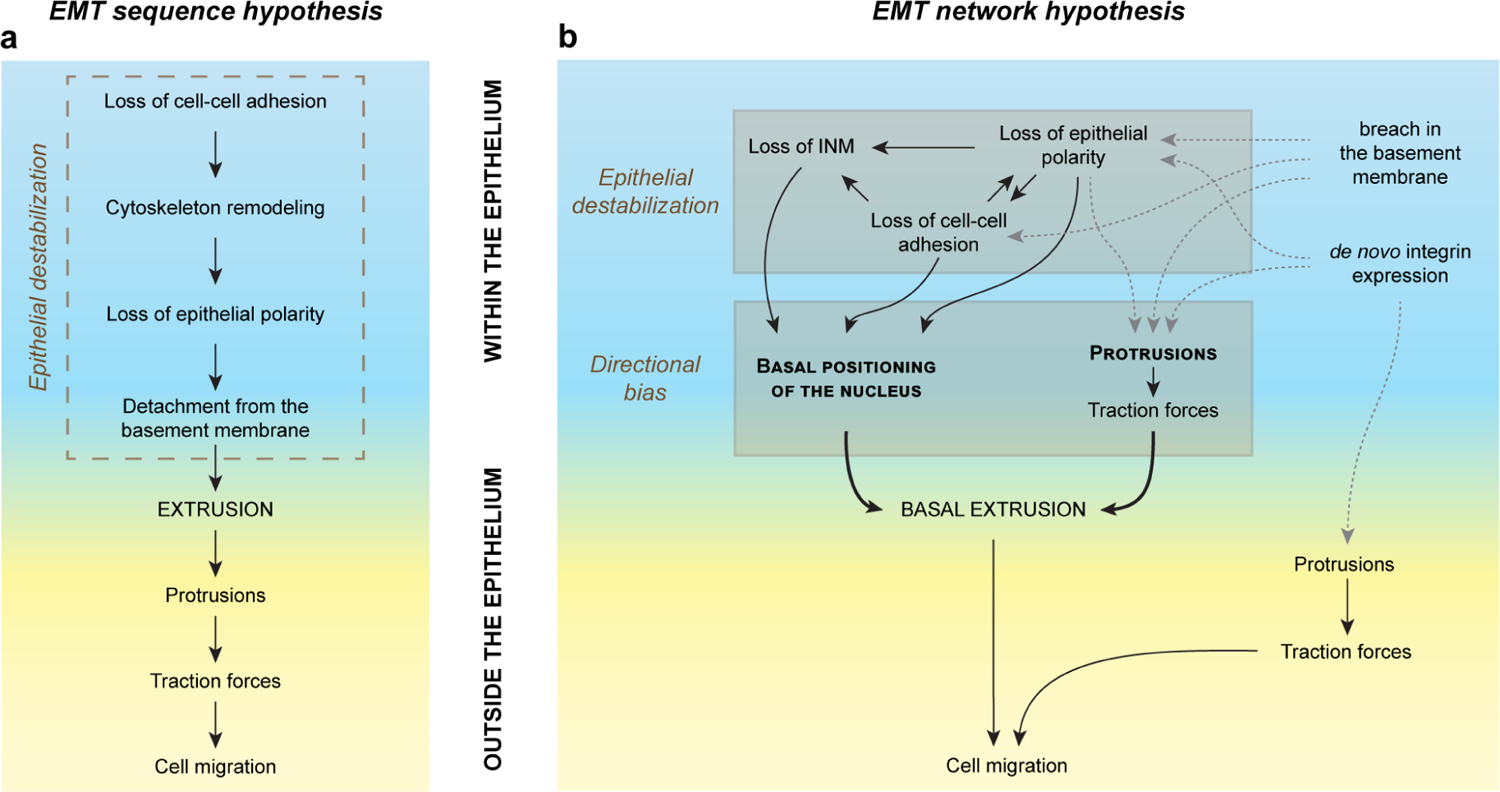
Timely and efficient basal extrusion downstream of EMT requires basal positioning of nuclei and protrusive activity a, Diagram representing the theoretical linear EMT cascade often used to describe EMT cell events in a logical manner. It starts with a module of epithelial destabilization leading to extrusion of cells into the extracellular matrix underlying the basal side of the tissue and ends with the adaptation of the cells to the local environment, adopting a migratory phenotype. Migration is only possible if extrusion occurs towards the basal side. An important limitation of this model is that epithelial destabilization in itself does not provide directionality of extrusion and can theoretically lead to either apical or basal extrusion. b, Diagram representing the alternative view of a non-linear array of EMT cellular events. The module of epithelial destabilization contains multiple interdependent events. This allows for multiple scenarios to co-exist to promote basal positioning of the nucleus, a key step favoring extrusion towards the basal side. In this model, loss of interkinetic movements and protrusive activity actively contribute to basal extrusion. This network of non-mandatory events supported by computational and biological data better fits the observed diversity of EMT scenarios, the documented heterogeneity of EMT cell populations and the current models of transcriptional regulation of EMT by a gene regulatory network.

## Discussion

Over the last two decades our view of EMT has dramatically evolved. Initially thought as a switch, then as a transition^2^, the various trajectories of cells to and from the mesenchymal state is now better encapsulated in the term epithelial-mesenchymal plasticity^51^. This change of perspective is due to multiple factors among which: i/ the understanding that the journey from E to M is not linear and reversible, ii/ the observations of many intermediate phenotypes in terms of gene expression signatures, iii/ the study of gene regulatory networks revealing complex interplay and feedback loops among upstream regulators of EMT. While the realization that cell populations undergoing EMT display diversity and heterogeneity took shape, our view of the molecular and cellular implementation of the conversion from E to M remained somewhat linear and thus at odds with the epithelial-mesenchymal plasticity model. The functional overlap between molecular players involved in cell polarity, adhesion and proliferation has dramatically hindered experimental progress on that front. The goal of our simulations was to conceptually fill the gap between the supposedly linear cascade of cell events and the observed diversity and heterogeneity of cell strategies in biological samples. Our results point to some observations that warrant further discussion.

The first one is about the impact of losing cell-cell adhesion (***A***). It is often presented as the first step of EMT because it makes sense that cells would detach from their neighbors to avoid “wasting” energy trying to pull on the extracellular matrix while being anchored in the epithelium. This notion is also supported by the early expression of numerous cadherin repressors such Snail1/2 and Twist in EMT cells^1-3, 10^. Yet, several ***in vivo*** examples show that loss of cadherins is neither a pre-requisite for other steps to follow suit nor sufficient to drive basal extrusion. Cephalic neural crest cell co-express E and N-cadherin at the time of EMT^19^. In addition, rather than promoting basal extrusion, interfering with cadherin levels in trunk neural crest cells randomizes extrusion leading some cells to fall into the lumen^52, 53^. During gastrulation of chicken and mouse embryos, cadherin-based junctions are conserved in ingressing cells^54-56^. Ingression is performed by a combination of degradation of the basal lamina and apical constriction^57, 58^, the latter requiring cell-cell adhesion. Similarly, in drosophila ventral furrow, modulation of cadherins is not a pre-requisite for ingression of the mesoderm^59^. Furthermore, in mammary gland epithelium maintenance of E-cadherin expression is required for Twist1-dependent basal extrusion^60^. Our simulations and experiments strongly indicate that epithelial destabilization in general leads to extrusion on both sides and that loss of apical adhesion (***A***) favors rapid and massive apical extrusion if not coupled to other events such as protrusions and/or loss of INM. The apical side is particularly prone to extrusion given the lack of physical obstacle outside of the tissue on this side compared to the basal side that is apposed against extracellular matrix. Thus, additional information atop epithelial destabilization is needed to generate a basal bias for extrusion towards the matrix. The examples described above and our simulations suggest that maintaining apical adhesion and its associated contractility might actively contribute to such needed bias.

Another interesting observation is the fact that interfering with the matrix and integrin expression lead to epithelial disorganization, moderate defect in cell polarity, non-apical mitoses and protrusive activity on both sides of the epithelium. It even led to some apical extrusion. This is at odds with the often proposed idea that a breach in the basal membrane may, in itself, be instructive for cells to leave the tissue. Performed independently, disorganizing laminin or overexpressing integrins are not sufficient to drive basal extrusion. Thus, what our data and simulations strongly suggest is the need for coordination between the breach in the basement membrane and other events such as protrusion as observed early ***in vivo*** in delaminating neural crest cells^20, 21^, apical constriction as observed in the primitive streak^56-58^ or basal positioning of the nucleus, either by loss of INM or synchrony with the S-phase^32, 33^, to ensure basal extrusion.

***In vivo*** observations of neural crest delamination^20, 21^ and our simulations point to an early role of protrusions. While this is in opposition with the logical view that extrusion of the cell body should precede activation of migratory behavior, it may make sense in the context of development. Neural crest cells must reach territories that are located far from their site of origin to form numerous structures such as cartilages and bones of the face or ganglia of the peripheral nervous system^10^. Therefore, activation of motility is a sign that cells are capable of embarking for the next step of their normal development. By contrast, failure to become motile in the context of epithelial destabilization might lead to apical extrusion and cell death. Thus acting as a selection mechanism.

The original complete EMT cascade involves loss of cell-cell adhesion and polarity and detachment from the basement membrane. These events lead to extrusion of the cell body and subsequent cell migration. One caveat of this cascade is that epithelial disorganization can lead to extrusion on either side, not specifically towards the basal compartment^61^. In addition, it does not fit with the aforementioned ***in vivo*** observations. These ***in vivo***data indicate that multiple cascades can co-exist. Overall, our study supports the notion that EMT is the result of an array of cell and molecular events among which prepositioning of the nucleus can be obtained through multiple inputs. That in itself opens the possibility of multiple scenarios. Therefore, it suggests that the observed heterogeneity is, at least in part, a by-product of the fact that successful basal extrusion is not linked to a single event. In addition to nuclear positioning, protrusive activity ensures timing and further imposes directionality and thus correct for other non-efficient scenarios that might otherwise lead to apical extrusion. Altogether, our conclusions help fill the gap between the expected diversity and heterogeneity in EMT cell populations from the EMT gene regulatory network and the actual implementation of cellular events. Furthermore, it strongly suggests that any event destabilizing epithelial organization might prime cells to extrude, in the context of primary tumors that would include hyper proliferation. Our study also strongly suggests that any molecular component of the motility machinery may be seen as an early rather than a late sign of putative successful basal extrusion and migration and, under some circumstances, invasion.

## Methods

### Chicken Eggs

Fertilized chicken eggs were purchased from S.C.A.L (Société commerciale avicole du Languedoc) and incubated at 38°C until the desired stage^62^.

### Electroporation

Embryos at stage HH12 were windowed. Using a glass capillary, a solution of 6% sucrose, 0.05% Fast Green containing the desired expression vectors is injected into the lumen of the posterior neural tube. A series of 7 square pulses of 80 ms and 5 Hz at 28V is delivered using a custom-made generator (CalTech workshop) and a pair of NepaGene electrodes (CUY611P7-4). A drop of Phosphate Buffer Saline (PBS) 1X is added before closing the egg and incubating it overnight. All expression vectors were used in a range of 1.5-2 µg/µL.

### Histology

After fixation in 4% paraformaldehyde, then prepared for sectioning with either the cryostat or the vibratome. For cryosections, embryos are washed in Phosphate Buffer (PB), incubated in PB / 15% sucrose overnight at 4°C. Embryos are transferred in PB / 15% sucrose / 7.5% gelatin (PBSG) for 2 hours at 42°C. Small weighting boats are used as molds. A thin layer of PBSG is deposited at the bottom and left to set. Embryos are transferred onto the layer using a plastic 2.5 mL pipette. Each embryo is placed in a drop of PBSG. When drops are set, the weighting boat is filled with PBSG and left to set on the bench. The dish is then placed at 4°C for 1 hour to harden the gelatin. Individual blocks are cut under a dissecting microscope to position the embryo in the desired orientation for sectioning. Blocks are frozen in isopentane (Sigma, 615838) at −70°C and stored at −70°C until sectioning. For vibratome sectioning, embryos are placed in PBS and transferred into to 5% low-melting agar in small petri dishes. Blocks are then cut with a scalpel and subsequently transferred in PBS for sectioning.

### Cryosections and immunostaining

Cryosections were performed using a cryostat Leica CM1950 as previously described^12^. Sections of 14 µm were incubated in PBS 1X for 30 minutes at 42°C to remove gelatin, treated with PBS 1X / 1% triton / 2% newborn calf serum for 1 hour for permeabilization and blocking. Vibratome 70 µm sections are obtained from Leica VT1000 S, then treated with 5%BSA / 1% triton for one hour at room temperature for permeabilization and blocking. Subsequent incubations with antibodies were performed under a coverslip for cryosections or floating for vibratome sections. Primary antibodies were diluted in PBS1X / 0.1% triton / 2% newborn calf serum for cryosections and 1% BSA / 0.1% Triton for vibratome sections. Sections were incubated with primary antibodies overnight at 4°C. Secondary antibodies were diluted in PBS 1X and applied on cryosections for 2 hours at room temperature or in 1% BSA / 0.1% Triton and incubated for 5h at room temperature for vibratome sections. All washes were done in PBS 1X. All antibodies were used at 1-2 µg/mL. Rabbit anti-snail2 (Cell Signaling, C19G7), rabbit anti-Sox9 (Millipore, AB5535), rabbit anti-PCM1^63^, mouse anti-phospho-histone 3 (Ser10) (Cell Signaling MA312B), mouse anti-aPKC (Santa Cruz, sc17781). The following antibodies, mouse anti-N-cadherin (DSHB, 6B3), mouse anti-Sox9 (DSHB, DA1D1), mouse anti-Fibronectin (DSHB, B3/D6), mouse anti-TFAP211(DSHB, 3B5), mouse anti-laminin (DSHB, 3H11), were obtained from the Developmental Studies Hybridoma Bank, created by the NICHD of the NIH and maintained at The University of Iowa, Department of Biology, Iowa City, IA 52242. Anti-mouse an anti-Rabbit secondary antibodies were coupled with alexa 647 and purchased from Invitrogen (Life Technology). For analysis of staining intensities on sections, nuclei were segmented using the surface tool in Imaris. Intensities in each channel were automatically retrieved and normalized to the peak value of each dataset.

### Image acquisition

Images were taken on a Zeiss 710 or a Leia SP8 confocal microscope. Images were then processed with FIJI or Imaris/BitPlane.

### Integrin expression vectors

The sequence corresponding to integrin α5 (Itga5) mRNA (GenBank ID: KC439457.1), including the 5’ UTR leader sequence and the complete coding sequence, fused in frame to a flexible linker^64^ and mCherry sequence was synthetized by GenScript and cloned using XbaI/XhoI sites in pCAGGS vector for electroporation (ITGA5:mCherry_pCAGGS). The complete coding sequence corresponding to the predicted integrin subunit α4 (Itga4) mRNA (NCBI Reference Sequence XM_040676057.2) was placed after the ITGA5 5’ UTR leader sequence, fused in frame to the flexible linker and eGFP sequence was synthetized by GenScript and cloned using XbaI/XhoI sites in pCAGGS vector for electroporation (ITGA4:EGFP_pCAGGS).

### Statistics

Statistical analyses were performed with Prism 6 (GraphPad). Datasets were tested for Gaussian distribution. Student t-test, or ANOVA followed by multiple comparisons were used with the appropriate parameters depending on the Gaussian vs non-Gaussian characteristics of the data distribution. Significance threshold was set at p < 0.05. Box and whiskers plot: the box extends from the 25th to the 75th percentile; the whiskers show the extent of the whole dataset. The median is plotted as a line inside the box. Point-Biserial correlation analyses between parameters and extrusion rates from simulations were performed as described in section 5.5 of the Supplementary Information.

### Enzymatic and drug treatments

10-somite long portions of the whole trunk at the level of the prospective forelimb region were dissected from embryos at stage HH12 as described previously^29, 38^. Explants were then cultured in suspension in DMEM for 2 hours at 37°C with a 1/250 dilution from 100 mM stock solution or Dispase II (Stem Cell Technologies; #07923, at 0.2 U/mL).

### Modeling

All details about the computational model are given in the Supplementary Information.

## Acknowledgements

We thank Dr. Andreas Merdès (CBI, CNRS UMR5077, Toulouse) for the kind gift of PCM1 antibody. ET is a research director at the French National Research Center (CNRS) and supported by grants from ANR SingleCrest (ANR-21-CE13-0028-02), the Association pour la Recherche contre le Cancer (ARC, ARCPJA22020060002084) and the Fondation pour la Recherche Médicale (FRM AJE201224). CD is a senior researcher at French National Institute for Medical Research (Inserm). BG is an assistant professor at Université de Toulouse. SMA’s research is funded by the Austrian Science Fund (FWF) through the project F65 and by the Vienna Science and Technology Fund (WWTF) [10.47379/VRG17014]. SP’s research was funded by the Vienna Science and Technology Fund (WWTF) [10.47379/VRG17014]. MAF’s research was funded by the Atmospheric Mathematics (AtMath) collaboration of the Faculty of Science of University of Helsinki, by the Academy of Finland, via an Academy project (project No. 339228) and a Finnish center of excellence (project No. 346306), and by the Centre for Mathematics of the University of Coimbra - UIDB/00324/2020 (funded by the Portuguese Government through FCT/MCTES). MAF was also partially supported by the Vienna Science and Technology Fund (WWTF) [10.47379/VRG17014]. We are grateful to Prof. Pierre Degond and Dr. Diane Peurichard for fruitful discussions throughout the project. We thank Drs. Benjamin Steventon (Cambridge University) and Valérie Lobjois (CBI, Toulouse) for critical reading of the manuscript.

## Author contributions

SP, MAF and SMA developed the computational model. SP and ET designed, performed and analyzed simulations. SP developed the stand-alone EMT simulator. CD, BG and ET designed, performed and analyzed the biological experiments. ET organized the data, prepared figures and wrote the manuscript. All authors commented on the manuscript.

