## Supplementary Figures for "Modelling variability and heterogeneity of EMT scenarios highlights nuclear positioning and protrusions as main drivers of extrusion"

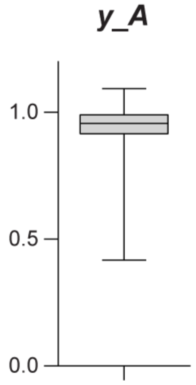


Figure S1. Normalized position along the apicobasal axis of nuclei at the time of *A* (*y_A*) in cells that performed apical extrusion from simulations shown in Figure 2a-c.

We performed 1000 simulations with one EMT-like cell implementing loss of cell-cell-adhesion (*A*) and retrieved the position of nuclei at time of *A* (*y_A*). The mean (0.94) and median (0.96) of this distribution are significantly different than the theoretical mean (0.5) and median (0.5) of a random distribution along the apicobasal axis. Mean comparison with one-sample t-test ($p < 0.0001$) and median comparison using Wilcoxon Signed Rank Test ($p < 0.0001$). This indicates that apically extruding cells at the time of *A* have their nuclei specifically located on the apical side of the tissue.

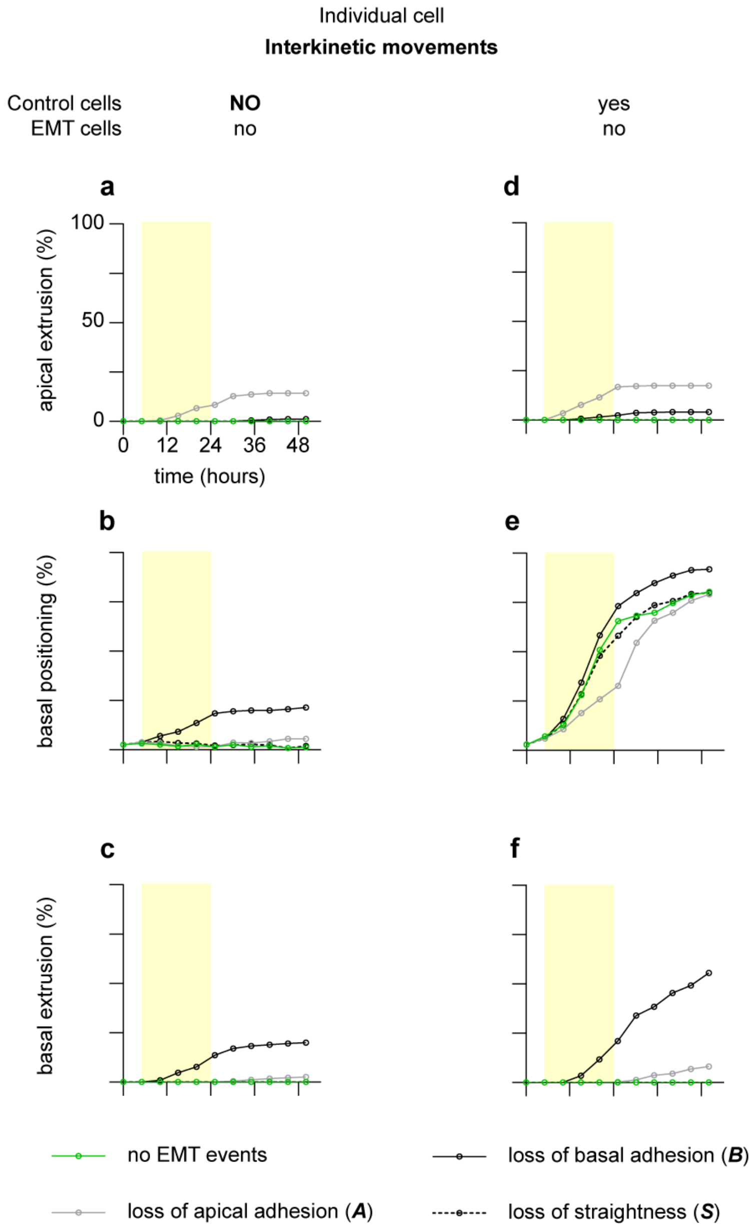


Figure S2. Cancelling interkinetic movements (INM) in control cells severely reduces basal positioning of EMT-like cells.

a-c, Rates of apical extrusion (a), basal positioning (b) and basal extrusion (c) of EMT-like cells from 500 simulations with one EMT-like cell without INM surrounded by controls without INM. d-e, Rates of apical extrusion (a), basal positioning (b) and basal extrusion (c) of EMT-like cells from 500 simulations with one EMT-like cell without INM surrounded by controls with INM. Panels d-e are identical to panels g-I from Figure 2 and are reproduced here to facilitate visual comparison with panels a-c. The yellow area corresponds to the time window of opportunity for EMT-like events to occur.

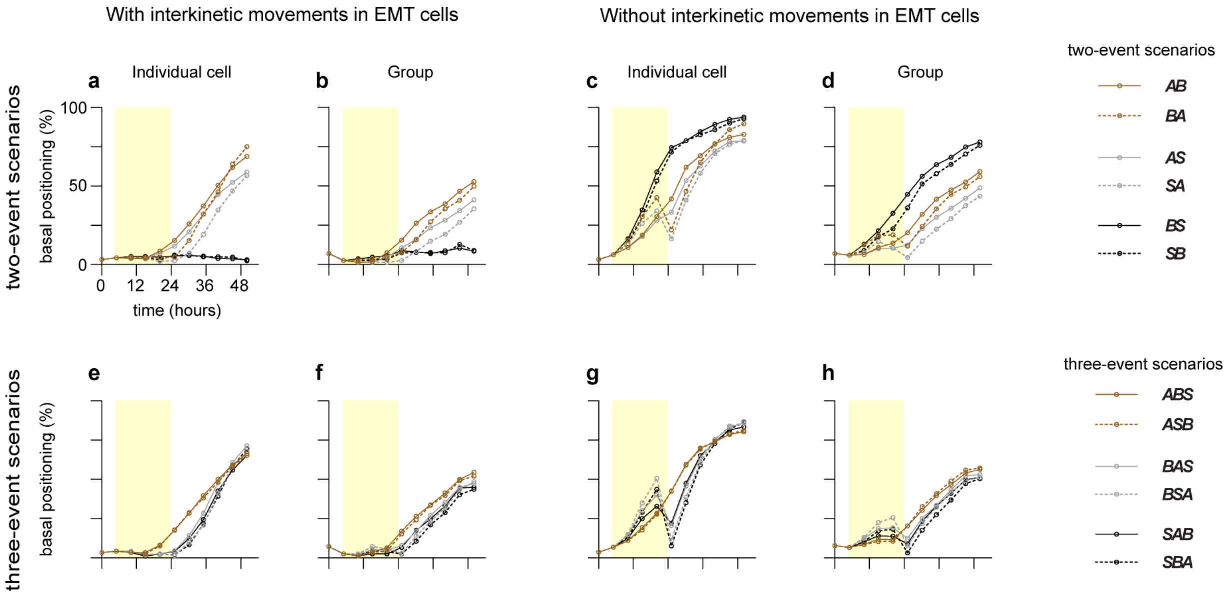


Figure S3. Rates of basal positioning of EMT cells two-event and three-event EMT-like scenarios.
**a-d**, Rates of basal positioning of EMT cells in two-event scenarios, with INM (a-b) or without INM (c-d), either individually (a, c) or in groups (b, d). **e-h**, Rates of basal positioning of EMT cells in three-event scenarios, with INM (e-f) or without INM (g-h), either individually (e, g) or in groups (f, h). These data are from simulations presented in Figures 3 and 4. The yellow area corresponds to the time window of opportunity for EMT-like events to occur.

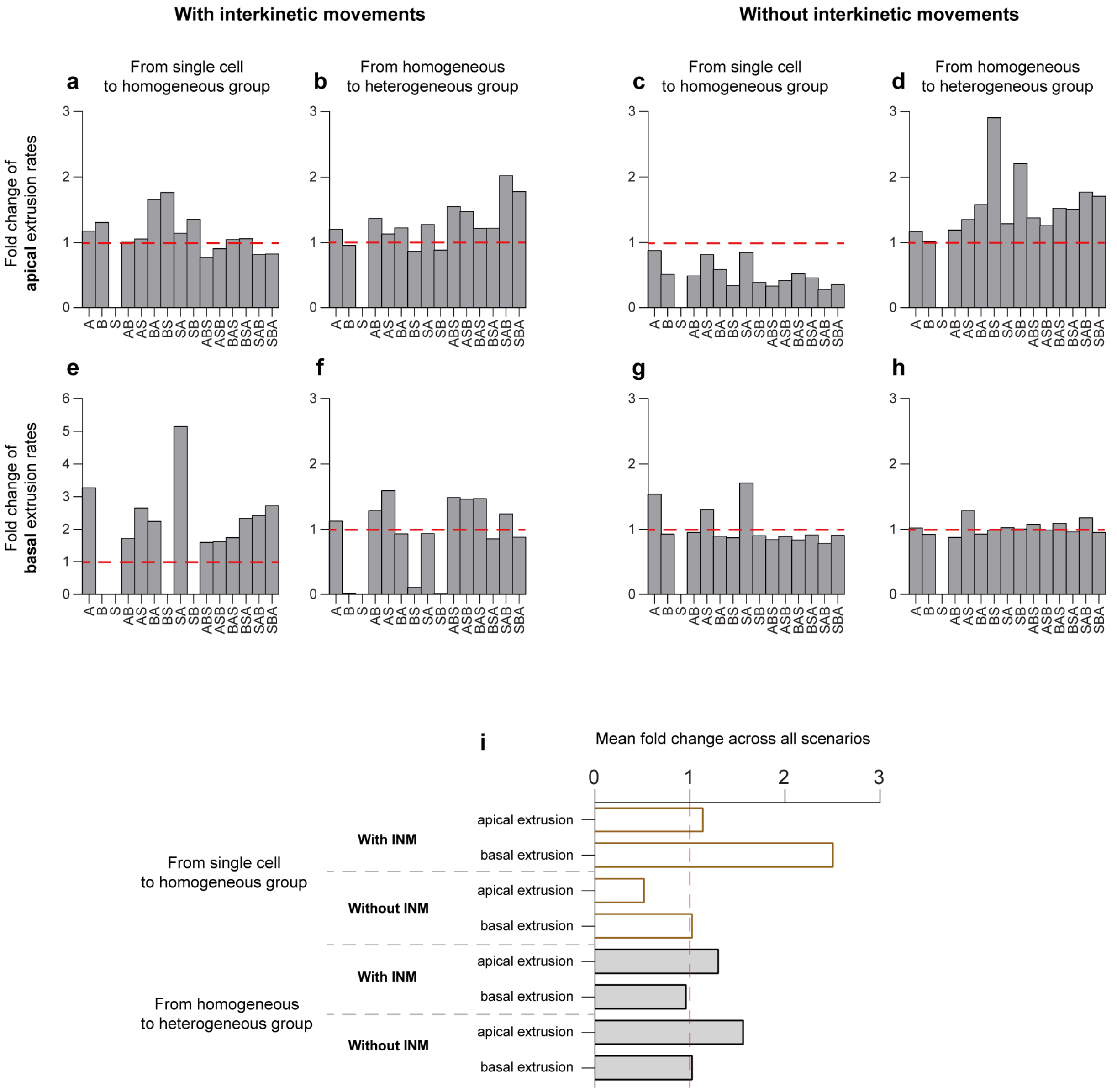


Figure S4. Fold change of apical and basal extrusion rates between simulations with individual cells, homogeneous groups and heterogeneous groups.

a-d, Fold change for apical extrusion rates with INM from single cell to homogeneous groups (a), from homogeneous to heterogeneous groups (b), without INM from single cell to homogeneous groups (c), from homogeneous to heterogeneous groups (d). e-h, fold change for basal extrusion rates with INM from single cell to homogeneous groups (e), from homogeneous to heterogeneous groups (f), without INM from single cell to homogeneous groups (g), from homogeneous to heterogeneous groups (h). i, Summary graph of mean fold per condition across all scenarios.

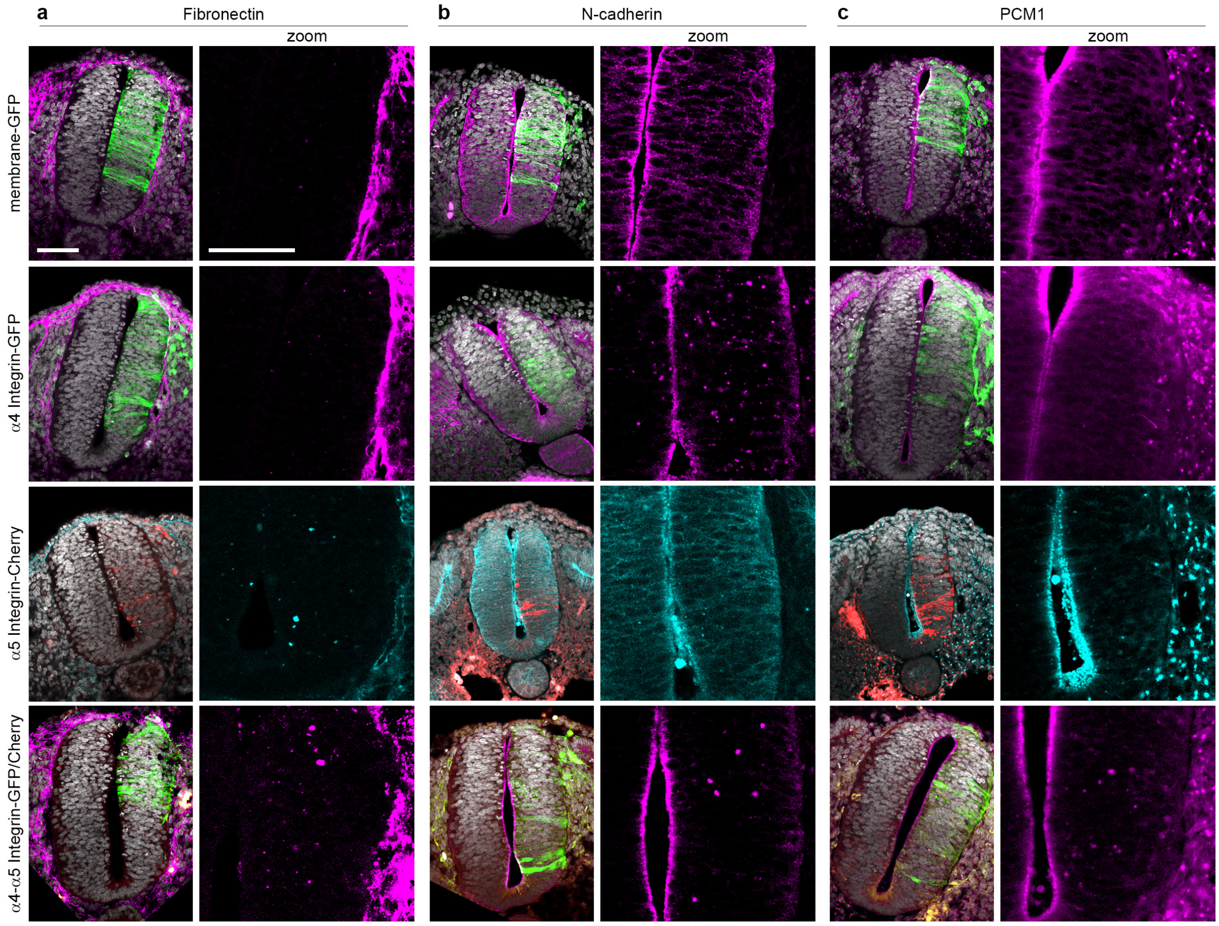


### Figure S5. Overexpression of α4 and α5 Integrins lead to moderate apicobasal defects

**a-c**, Representative images for immunostaining on cryosections against atypical Fibronectin (a; embryos $n_{mbGFP} = 3$, $n_{\alpha4} = 7$, $n_{\alpha5} = 2$, $n_{\alpha4+\alpha5} = 4$ ), N-cadherin (b; embryos $n_{mbGFP} = 3$, $n_{\alpha4} = 12$, $n_{\alpha5} = 6$, $n_{\alpha4+\alpha5} = 11$) and Pericentriolar Material 1 (PCM1, c; embryos $n_{mbGFP} = 3$, $n_{\alpha4} = 13$, $n_{\alpha5} = 6$, $n_{\alpha4+\alpha5} = 6$) from embryos expressing membrane-GFP, α4-Integrin, α5-Integrin or a combination of α4 and α5 Integrins, as indicated. α4 and α5 are displayed in green and red respectively. Immunostainings are shown in magenta for membrane-GFP, α4 and the α4 plus α5 conditions or in cyan for the α5 condition. Scale bar 80 µm. Images for other apical/basal markers and quantifications are shown in Figure 8.

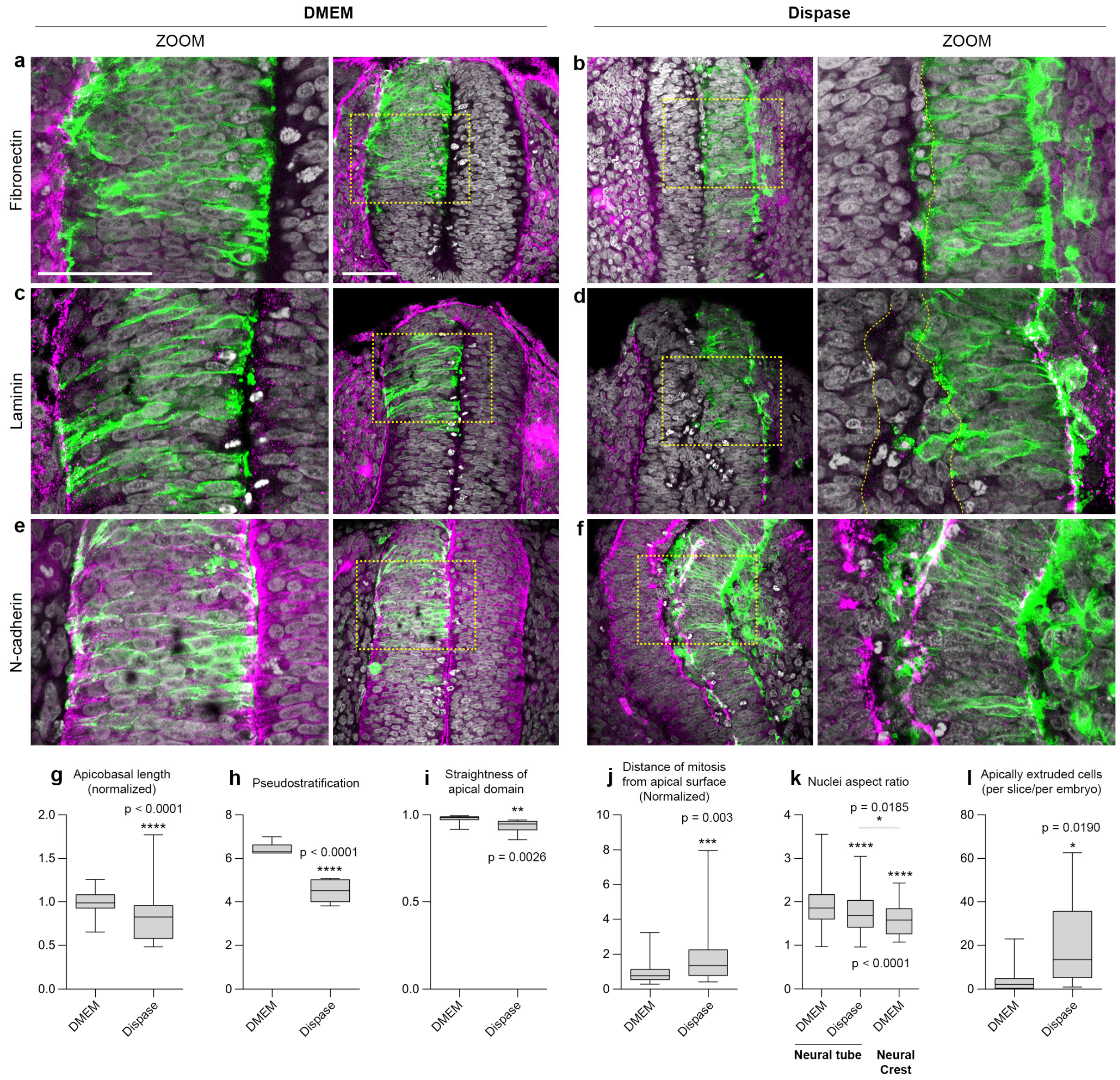


### Figure S6. Short-term destabilization of the basement membrane via enzymatic degradation

**a-f**, Immunostaining on cryosections from caudal parts of chicken embryos incubated with control culture medium (DMEM) or DMEM containing Dispase II (0.2 U/mL) for 2 hours. Sections were stained with anti-Fibronectin (**a-b**), anti-Laminin (**c-d**) or anti-N-cadherin (**e-f**). Yellow squares indicate the position of the zooms. Yellow dotted lines in panels b and d mark the position of the apical domain. Scale bars, 80 µm on low magnifications, 50 µm on zooms. **g**, Normalized apicobasal length (DMEM, n = 54 measurements averaged per embryo ; Dispase, n = 69 measurements averaged per embryo; Kolmogorov-Smirnov test). **h**, Pseudostratification, mean number of layers of nuclei along the apicobasal axis (DMEM, n = 96, averaged per embryo) ; Dispase, n = 112 measurements averaged per embryo; Unpaired t test with Welch's correction). **i**, Straightness of the apical domain (DMEM, n = 12; Dispase, n = 12; Unpaired t test with Welch's correction). **j**, Normalized distance of mitosis from apical domain (DMEM, n = 46; Dispase, n = 55; Mann Whitney test). **k**, Nuclei aspect ratio (Neural tube: DMEM, n = 207; Dispase, n = 202; Neural crest: DMEM, n = 75; one-way ANOVA, uncorrected Fisher's LSD). **l**, Mean number of apically extruded cells averaged per section per embryo (DMEM, n = 10; Dispase, n = 6; Kolmogorov-Smirnov test). Scale bars, 50 µm.
