## Supplementary material for "Modelling variability and heterogeneity of EMT scenarios highlights nuclear positioning and protrusions as main drivers of extrusion": S1_Table

| Figure | Number of EMT-like cells | INM in EMT-like cells | Protrusions in EMT-like cells | EMT scenario | Heterogeneity of EMT cell clusters |
|---|---|---|---|---|---|
| Figure 2a-c | 1 | **ON** | OFF | A, B or S | OFF |
| Figure 2d-f | 11 | **ON** | OFF | A, B or S | OFF |
| Figure 2g-i | 1 | OFF | OFF | A, B or S | OFF |
| Figure 2j-l | 11 | OFF | OFF | A, B or S | OFF |
| Figure 3a-b | 1 | **ON** | OFF | AS, SA, BS, SB, AB, or BA | OFF |
| Figure 3c-d | 11 | **ON** | OFF | AS, SA, BS, SB, AB, or BA | OFF |
| Figure 3e-f | 1 | OFF | OFF | AS, SA, BS, SB, AB, or BA | OFF |
| Figure 3g-h | 11 | OFF | OFF | AS, SA, BS, SB, AB, or BA | OFF |
| Figure 4a-b | 1 | **ON** | OFF | ABS, ASB, BAS, BSA, SAB or SBA | OFF |
| Figure 4c-d | 11 | **ON** | OFF | ABS, ASB, BAS, BSA, SAB or SBA | OFF |
| Figure 4e-f | 1 | OFF | OFF | ABS, ASB, BAS, BSA, SAB or SBA | OFF |
| Figure 4g-h | 11 | OFF | OFF | ABS, ASB, BAS, BSA, SAB or SBA | OFF |
| Figure 5a-b (individual cells INM) | 1 | **ON** | OFF | A, B, S, AB, AS, BA, BS, SA, SB, ABS, ASB, BAS, BSA, SAB or SBA | OFF |
| Figure 5a-b (individual cells no INM) | 1 | OFF | OFF | A, B, S, AB, AS, BA, BS, SA, SB, ABS, ASB, BAS, BSA, SAB or SBA | OFF |
| Figure 5a-b (homogeneous groups) | 11 | **ON** or OFF | OFF | A, B, S, AB, AS, BA, BS, SA, SB, ABS, ASB, BAS, BSA, SAB or SBA | OFF |
| Figure 5a-b (Heterogeneous groups) | 11 | **ON** or OFF | OFF | Chosen at random per cell in the group | **ON** |
| Figure 5e-f Heterogeneous groups INM | 11 | **ON** | OFF | Chosen at random per cell in the group | **ON** |
| Figure 5e-f Heterogeneous groups no INM | 11 | **OFF** | OFF | Chosen at random per cell in the group | **ON** |
| Figure 5e-f Heterogeneous groups 50% INM | 11 | **50%** | OFF | Chosen at random per cell in the group | **ON** |
| Figure 5e-f, Figure 6 Heterogeneous groups 50% INM, 50% ***P*** | 11 | **50%** | 50% | Chosen at random per cell in the group | **ON** |

**Supplementary Table 1. Combinations of parameters to perform simulations described in the Figures and Supplementary Figures using sEMTor.** https://semtor.github.io/
