## Supplementary Information for "Modelling variability and heterogeneity of EMT scenarios highlights nuclear positioning and protrusions as main drivers of extrusion"

### Agent-based model for epithelial-to-mesenchymal transitions

#### 1 Overview

##### 1.1 Description of the pseudostratified epithelium (PSE) model

We use as a biological reference for PSE the neuroepithelium of the chicken embryo at two days of development. We consider as the main mechanical components the cell nucleus, the cell's cytoskeleton and the points of adhesion between cells at their apical point and between the cell's basal point and the underlying basal layer of the tissue. Given that in PSE, the cells are elongated along the apicobasal axis and that the largest physical object is the nucleus, we simplify the interaction between cells by mainly considering interaction between nuclei. Other parameters, such as the distance between the basal points of neighboring cells, can further account for lateral cell-cell interaction.

As governing equations, we use an overdamped Newton equation with inequality constraints to model cell volume exclusion. Some parameters change depending on the cell cycle.

Our PSE model is equivalent to the model in [1] with the new additions of EMT-like events and a new mathematical formulation with improved numerical stability.

The model incorporates the following elements:

- **Volume exclusion and elasticity of nuclei.** The mechanical pushing between nuclei leads to the pseudostratified nature of the tissue. As such, it is important to model volume exclusion. However, the nuclei are not just rigid objects, instead, they can deform, that allows them to squeeze through small spaces in crowded environments. To capture the volume exclusion and the non-rigid nature, we model each nucleus as a set of two spheres. An inner hard sphere with strict

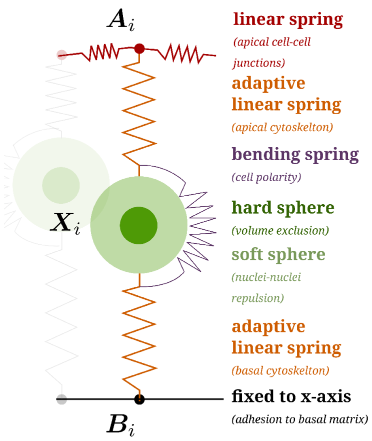


Figure 1: Overview of the model components for a single cell. The green spheres represent the cell nucleus and the adaptive springs model the cytoskeleton. The apical network consists of linear springs connecting neighbouring points. The basal network is a static line to which the basal points of the cells are constrained. Interkinetic nuclear movement is modelled by changing the rest lengths of the adaptive springs.

non-overlap conditions to other nuclei. We complement this with an outer soft sphere with repulsive forces between with soft spheres of other nuclei.

- **Apical cell-cell junctions.** The apical side of the neural tube is at the boundary to the central lumen. The apical side does not provide any organised structure for cells to grab onto. Instead, the cells themselves form the apical layer of the tissue, which consists of a network of cell junctions with adhesive contacts. For a simple two-dimensional model of the tissue, we simplify this structure to a line of apical points which represent the cell position at the apical layer. These apical points are connected with linear springs to their neighbours, which models the adhesive bonds.

- **Basal cell-matrix adhesion.** The situation is different on the basal side where cells are anchored to the extracellular matrix. In the model, we introduce basal points which represent the most basal part of each cell which is connected to the matrix. We model the basal matrix as a static line with the condition that basal points have to lie on the fixed basal line.

- **Cytoskeleton.** The cytoskeleton regulates the position of the cell nuclei via microtubules. We model the cytoskeleton as two linear springs that span between the basal point and the nuclei center, and the apical point and the nuclei center.

- **Interkinetic nuclear movement (INM).** During proliferation, cells progress through the various phases of cell cycle. In the G1 phase, S phase and early (passive) G2 phase of the cell cycle, the rest lengths of these cytoskeleton springs will change in a way to match the current extension of the cytoskeleton. During the late (active) G2 phase and mitosis, the nuclei move apically. This rapid apical movement is modelled by changing the rest lengths of the basal- nuclei and apical-nuclei springs such that the most apical position is the equilibrium for both springs.

- **Proliferation.** At the end of the cell cycle, cells will divide into two daughter cells. However, since we only model a 2D slice of the tissue, we have to account for the fact that in the real tissue, not all daughters end up in the same 2D slice. Therefore, only in 20 % of all cell divisions a second daughter is created within the simulated 2D slice. To account for the increased stiffness of cells during mitosis, we also increase the radius of the hard sphere during the mitotic phase.

Putting all these rules together, we obtain a model of growing epithelial tissue. The growth is created by INM, proliferation and the combination of the right stiffness coefficients for apical-apical junctions and cytoskeleton springs.

The mathematical details are described in the next section.

##### 1.2 Description of the EMT model

Our EMT model extends the PSE model [1] by introducing a second cell type, which we refer to as **EMT cells**. These cells are initially of epithelial phenotype, but each cell has random time points at which it can perform a selection of the four EMT-like events $\boldsymbol{A}$, $\boldsymbol{B}$, $\boldsymbol{S}$ and $\boldsymbol{P}$. These events represent the loss of apical adhesion, loss of basal adhesion, loss of straigthness (alignment between the apical point, the center of the nucleus and the basal point) and protrusive activity. These events are described in detail below:

- **$\boldsymbol{A}$: Loss of apical adhesion.** When cells lose cell-cell adhesion on the apical side, then the cell's cytoskeleton will contract and the surrounding epithelial cells create new cell junctions to fill the gap in the apical layer. In our model, we implement these steps by removing the apical point from the apical network, setting the desired rest length of the apical-nuclei spring to zero and creating a new linear spring between apical points of the adjacent epithelial cells.
- **$\boldsymbol{B}$: Loss of basal adhesion.** Loss of adhesion to the basal membrane is modelled by removing the basal point from the basal layer and setting the desired rest length of the basal-nuclei spring to zero.
- **$\boldsymbol{S}$: Relaxation of cell straightness.** Epithelial cells have an apical-basal polarity that can be completely or partially lost during EMT. In our model the polarity is indirectly represented by the apical and the basal points which are at opposite sides of the nucleus due to the bending spring. We model the relaxation of straightness by decreasing the stiffness of the bending spring, which has the effect that the apical point, nucleus and basal points might no longer be in that order and/or not located on a straight line. This can potentially represents multiple events, a (partial) loss of polarity, a change of cell shape, as one can extrapolate the boundaries of the

actual cells from the relative positions of the apical/basal points and the nucleus but also a softening of the cells since cells with a weak bending spring will offer less resistance to local pressure coming from neighboring cells.

- $\boldsymbol{P}$**: Protrusions and active motion.** The area outside the basal region of the epithelium is called the extracellular matrix (ECM), which is a network of fibers of various proteins such as collagens. In the model, we do not represent fibers of the extracellular matrix. However, after event $\boldsymbol{B}$, a cell can extend its basal spring and attach its basal point to a new location located below the basal line and pull on it. If the basal point is already detached from the basal membrane, cells can protrude by moving their basal point in the direction of the basal-nuclei spring and as soon as the basal point is below the basal layer, we set the desired rest length of the basal-nuclei spring to zero. This seek-and-grab behavior is akin to a protrusive behavior. Note that $\boldsymbol{P}$ is a secondary EMT-like event and can only occur if $\boldsymbol{B}$ occurred.

We also are considering the following changes:

- **Stop of interkinetic nuclear movement (INM).** In some simulations, we selected that EMT cells might not perform interkinetic nuclear movement (INM) from the beginning of the simulation.
- **Heterogeneity of EMT.** The timing of the EMT-like events and the possibility for inhibition of INM are the source of heterogeneity in EMT in our model, in particular:
  - Not all EMT cells undergo all events $\boldsymbol{A}$, $\boldsymbol{B}$, $\boldsymbol{S}$, $\boldsymbol{P}$ and stop of INM.
  - EMT events happen in different sequential order and at different times.
  - Epithelial cells can become EMT cells in isolation or as a cluster.

These extensions of the PSE model form the foundation for our EMT model. We describe the mathematical details next.

##### 1.3 Online simulation: sEMTor – The EMT simulator

The model described in this document is accessible as a web-based, stand-alone EMT simulator: `https://semtor.github.io`. The simulation does not require any setup and allows trying different setups of the model, ranging from single individual EMT-like events to simulations with multiple cells and heterogeneous EMT.

### 2 Mathematical model

#### 2.1 Notation

In the following we will use the notation $\nabla W(x) := (\mathrm{D}W(x))^T$ to denote the gradient as the transpose of Jacobian of a function $W : \mathbb{R}^d \to \mathbb{R}$. We denote a collection of positions as $(x_1, \dots, x_N) \in \mathbb{R}^{2N}$ and we refer to the $y$ coordinate of a position via $y(x_i)$ where $y((a,b)^T) := b$. We denote the uniform distribution on an interval $[a, b]$ as $\mathcal{U}(a, b)$.

#### 2.2 The PSE model

Our PSE model is essentially equivalent to [1] but with the modification that we consider overdamped motion instead of a quasi-steady state simulation. This change does not impact the simulation outcome, but allows us to employ the position-based dynamics [2, 3] method, which leads to a sizeable runtime improvement. The numerical method is described in Section 4.

We consider a tissue with $N$ cells where the $i$th cell has the apical point as position $a_i \in \mathbb{R}^2$, the basal point at $b_i \in \mathbb{R}^2$ and the center of the nuclei is at $x_i \in \mathbb{R}^2$. We write $\boldsymbol{z} = (\boldsymbol{a}, \boldsymbol{x}, \boldsymbol{b}) = (a_1, \dots, x_1, \dots, b_1, \dots) \in \mathbb{R}^{6N}$ to denote all positions at once.

The mechanical description of the cell dynamics is given by the potential energy $W$, and $m \in \mathbb{N}$ many inequality constraints $g_1, \dots, g_m : \mathbb{R}^{6N} \to \mathbb{R}$ which implement for example non-overlap conditions.

As dynamic law, we use an overdamped Newton equation with Lagrangian multipliers for the inequality conditions. This yields a dynamic complementarity system

$$\frac{\mathrm{d}\boldsymbol{z}(t)}{\mathrm{d}t} = -\nabla_{\boldsymbol{z}} W(\boldsymbol{z}(t)) - \sum_{j=1}^{m} \lambda_j \nabla_{\boldsymbol{z}} g_j(\boldsymbol{z}) \tag{1}$$

$$g_j(\boldsymbol{z}) \geq 0, \quad \lambda_j \geq 0, \quad \text{and} \quad \lambda_j g_j(\boldsymbol{z}) = 0 \tag{2}$$

where $\lambda_1, \dots, \lambda_m \in \mathbb{R}^d$ denotes the Lagrangian multipliers which are subject to the Signorini conditions (2).

We note that we also include an additive noise term for the final equations, which is not included in (1) and (2) but explained later in Section 2.2.8.

Next, we will describe the different components of the model in the following order:

1. Cell cycle and age of cells,
2. Cyctoskeleton model,
3. Interkinetic nuclear movement (INM),
4. Apical cell junctions,
5. Cell division,
6. Forces in the system (energy terms),
7. Inequality constraints,

8. Noise,
9. Initial conditions.

##### 2.2.1 Cell cycle and age of cells

Each cell has a time of birth $t_i^{\text{birth}}$ and an age $t_i^{\text{age}} = t - t_i^{\text{birth}}$.

At the birth of a new cell, the duration of the cell cycle is randomly assigned as

$$t_i^{\text{cycle}} \sim \mathcal{U}(T^{\text{min-age}}, T^{\text{max-age}}), \tag{3}$$

where $T^{\text{min-age}} < T^{\text{max-age}}$ are the parameters for the minimal and maximal lifespan for each cell.

The cell cycle is divided into a passive phase (G1, S phase and early G2 phase), the G2 phase and mitosis. The parameters $T^{\text{G2}}$ and $T^{\text{mitosis}}$ denote the duration of these phases.

Accordingly, the mitosis phase of the $i$th cell starts when $t_i^{\text{age}} = t_i^{\text{cycle}} - T^{\text{mitosis}}$ and the G2 phase starts when $t_i^{\text{age}} = t_i^{\text{cycle}} - T^{\text{mitosis}} - T^{\text{G2}}$.

##### 2.2.2 Cytoskeleton model: adaptive rest lengths

The cytoskeleton is modelled by apical-nuclei and basal-nuclei springs. The apical and basal springs of the $i$th cell have adaptive rest lengths, which we denote by $\eta_i^{\text{ax}} \in \mathbb{R}$ and $\eta_i^{\text{bx}} \in \mathbb{R}$. The indices "ax", "bx" are used to denote values related to the apical-nuclei and basal-nuclei springs.

Moreover, we introduce the desired rest lengths $\eta_i^{\text{ax-des}}, \eta_i^{\text{bx-des}} \in \mathbb{R}$ which represent the direction into which the cell would like to move. The rest lengths update according to the following differential equations

$$\frac{\mathrm{d}\eta_i^{\text{ax}}}{\mathrm{d}t} = -k^{\text{ax}}(\eta_i^{\text{ax}} - \eta_i^{\text{ax-des}}), \tag{4}$$

$$\frac{\mathrm{d}\eta_i^{\text{bx}}}{\mathrm{d}t} = -k^{\text{bx}}(\eta_i^{\text{bx}} - \eta_i^{\text{bx-des}}). \tag{5}$$

For most of the cell cycle, the cells are passive ($t_i^{\text{age}} < t_i^{\text{cycle}} - T^{\text{G2}} - T^{\text{mitosis}}$) which we model with the choice

$$\eta_i^{\text{ax-des}} = \|x_i - a_i\| - R^{\text{soft}}, \tag{6}$$

$$\eta_i^{\text{bx-des}} = \|x_i - b_i\| - R^{\text{soft}}. \tag{7}$$

###### 2.2.3 Interkinetic nuclear migration (INM)

The nuclei movement during the cell cycle is controlled by changing the desired rest lengths of the cells' cytoskeletons.

At the onset of the active G2 phase, the apical cytoskeleton contracts and the basal cytoskeleton extends, which leads to an upward migration of the cells to prepare for cell division. Therefore, when $t_i^{\text{age}} \geq t_i^{\text{cycle}} - T^{\text{G2}} - T^{\text{mitosis}}$, we set as

$$\eta_i^{\text{ax-des}} = 0 \tag{8}$$

$$\eta_i^{\text{bx-des}} = \|a_i - b_i\| - 2R^{\text{soft}}. \tag{9}$$

During the mitosis phase, the cell also extends its volume, which we model by increasing the parameter

$$R_i^{\text{hard}}(t) = R^{\text{hard,G2}} \tag{10}$$

when $t_i^{\text{age}} \geq t_i^{\text{cycle}} - T^{\text{mitosis}}$.

###### 2.2.4 Apical junctions

For apical junctions, we use linear springs with dynamic rest length, similar as for the cytoskeleton. Apical junctions exists between cells which are next to each other on the apical side of the tissue. Therefore, we define the edges of the apical layer as a set

$$\mathcal{A} = \{(1,2), (2,3), \ldots, (N-1, N)\}. \tag{11}$$

Each apical junction modelled as linear spring with rest length $\eta_{ij}^{\text{aa}}$ where $(i,j) \in \mathcal{A}$ are indices of apically adjacent cells.

The rest length of the apical-apical springs changes according to

$$\frac{\mathrm{d}\eta_{ij}^{\text{aa}}}{\mathrm{d}t} = -k^{\text{aa}}\eta_{ij}^{\text{aa}} \quad \text{for all } (i,j) \in \mathcal{A}, \tag{12}$$

where $k^{\text{aa}}$ denotes the speed of adaptation.

###### 2.2.5 Cell division

Since we only model a 2D slice of a three-dimensional tissue, the growth of the tissue would be too fast if cells divide at a realistic rate. Instead of modifying the cell cycle, we consider that with a probability

$$p^{\text{div}} \in (0,1)$$

both daughter cells will be part of the modelled 2D slice but in the other cases the second daughter cell will be "outside" of the simulation slice and hence no cell is added to the simulation.

**Cell division with two offsprings.** Let $t$ denote the time of cell division for the $i$th cell and $t^+$ denote the time right after the cell division occured. To implement cell division, we increase the number of cells $N \mapsto N+1$. The new offsprings have indices $i$ and $N+1$. We define the offset

$$\Delta X = 0.05\mu\text{m} \begin{pmatrix} 1 \\ 0 \end{pmatrix},$$

which determines the new positions of the two offsprings as

$$x_i(t^+) = x_i(t) - \Delta X, \quad x_{N+1}(t^+) = x_i(t) - \Delta X, \tag{13}$$

$$a_i(t^+) = a_i(t) - \Delta X, \quad a_{N+1}(t^+) = a_i(t) - \Delta X, \tag{14}$$

$$b_i(t^+) = b_i(t) - \Delta X, \quad b_{N+1}(t^+) = b_i(t) - \Delta X. \tag{15}$$

Correspondingly, we remodel the apical network by adding the edge $(i, N+1)$ and by connecting the former right neighbour of the $i$th cell with the new $N+1$th cell. In the same way, we remodel the basal network (which will be defined later in Section 2.2.7).

For both cells, we revert the increase of the hard radius (10), i.e. we set $R_i^{\text{hard}}(t^+) = R^{\text{hard}}$ and $R_{N+1}^{\text{hard}}(t^+) = R^{\text{hard}}$.

Finally, we update the cell cycle duration and the birth times:

$$t_i^{\text{cycle}}, t_{N+1}^{\text{cycle}} \sim \mathcal{U}(T^{\text{min-age}}, T^{\text{max-age}}),$$
$$t_{N+1}^{\text{birth}} = t_{N+1}^{\text{birth}} = t.$$

**Division with one offspring.** Here, we do not spaw a new cell and just update the hard radius, cell cycle duration and cell birth time for the $i$th cell exactly as in the two offspring case.

##### 2.2.6 Energy terms and forces

The potential energy of the system is composed by the two linear springs (apical and basal cytoskeleton), linear springs connecting adjacent apical points, the bending spring (cell straightness) and finally the potential for pair-wise soft-repulsion between nuclei.

We define the energy terms in their non-dimensional form as in Ferreira *et al.* [1].

The apical and basal linear springs have the potential energy

$$W_i^{\text{ax}} = \frac{\alpha^{\text{ax}}}{R^{\text{soft}} + \eta_i^{\text{ax}}} \left( \|x_i - a_i\| - R^{\text{soft}} - \eta_i^{\text{ax}} \right)^2, \tag{16}$$

$$W_i^{\text{bx}} = \frac{\alpha^{\text{bx}}}{R^{\text{soft}} + \eta_i^{\text{bx}}} \left( \|x_i - b_i\| - R^{\text{soft}} - \eta_i^{\text{bx}} \right)^2 \tag{17}$$

where $\alpha^{\text{ax}}, \alpha^{\text{bx}}$ are the stiffness parameters for the apical and basal cytoskeletons, respectively. The radius $R_i^{\text{soft}}$ denotes the radius of soft replulsion of each cell. We recall that $\eta_i^{\text{ax}}, \eta_i^{\text{bx}}$ are the (dynamic) rest lengths of the apical and basal cytoskeletons.

The apical-apical cell junctions are also linear springs

$$W_{ij}^{\text{aa}} = \frac{\alpha^{\text{aa}}}{2R^{\text{soft}}} \left( \|a_i - a_j\| - \eta_{ij}^{\text{aa}} \right)^2 \quad \text{for all } (i,j) \in \mathcal{A},$$

where $\alpha^{\text{aa}}$ denotes the stiffness coefficient and $\eta_{ij}^{\text{aa}}$ is the rest length of the apical-apical springs.

The soft repulsion between nuclei is modelled by the potential energy

$$W_{ij}^{\text{xx}} = \begin{cases} \frac{\alpha^{\text{xx}}}{2R^{\text{soft}}} \left( \|x_i - x_j\| - R^{\text{soft}} \right)^2 & \text{if } \|x_i - x_j\| < 2R^{\text{soft}}, \\ 0 & \text{otherwise} \end{cases}$$

where $\alpha^{\text{xx}}$ denotes the repulsion stiffness and $R^{\text{soft}}$ is the radius of soft repulsion.

Finally, the straigthness energy term has the form of a bending spring which depends on the angle between the apical-nuclei and the basal-nuclei cytoskeleton

$$W_i^{\text{axb}} = \alpha^{\text{axb}} \left( \cos(\theta_i) + 1 \right)^2$$

where $\alpha^{\text{axb}}$ denotes the stiffness coefficient and $\theta_i$ is the angle between the apical and basal cytoskeletons of the $i$th cell, i.e.

$$\cos(\theta_i) = \frac{(a_i - x_i) \cdot (b_i - x_i)}{\|a_i - x_i\| \|b_i - x_i\|}.$$

The total potential energy is given by

$$W = \sum_{i=1}^{N} \left( W_i^{\text{ax}} + W_i^{\text{bx}} + W_i^{\text{axb}} \right) + \sum_{(i,j) \in \mathcal{A}} W_{ij}^{\text{aa}} + \sum_{i=1}^{N} \sum_{j=1}^{i-1} W_{ij}^{\text{xx}}.$$

##### 2.2.7 Constraints

We consider four kinds of constraints to implement volume exclusion between cells and the dynamics at the basal membrane.

Volume exclusion leads to a strict non-overlap constraint between the hard spheres of each cell, i.e.,

$$\|x_j - x_i\| \geq R_i^{\text{hard}} + R_j^{\text{hard}} \quad \text{for all } 1 \leq i < j \leq N \tag{18}$$

where $R_i^{\text{hard}}$ denotes the hard radius of the $i$th cell.

The other constraints determine the restriction of the basal points $b_i$ to the basal layer. We consider the basal layer as relatively static compared to the apical layer. Therefore, we restrain the basal points to the $x$-axis. We recall that we denote the $y$-coordinate of the basal points by $y(b_i)$, then, the constraint reads

$$y(b_i) = 0 \quad \text{for all } 1 \leq i \leq N. \tag{19}$$

Effectively, this constraint reduces the degrees of freedom, but this constraint becomes inactive during EMT, see Section 2.3.2.

Similar to the apical network, we also consider the basal network as a set of edges which is initially

$$\mathcal{B} = \{(1,2), (2,3), \ldots, (N-1, N)\}. \tag{20}$$

The third constraint enforces that the order of basal points remains invariant. Therefore, we require

$$y(b_i) \leq y(b_j) \quad \text{for all } (i,j) \in \mathcal{A}. \tag{21}$$

The last constraint introduces a maximal distance $d^{\text{max-basal}}$ between connected basal points,

$$\|b_i - b_j\| \leq d^{\text{max-basal}} \quad \text{for all } (i,j) \in \mathcal{B}. \tag{22}$$

Finally, we define the constraint functions $g_1, \ldots, g_m : \mathbb{R}^{6N} \to \mathbb{R}$ for $m = \frac{N(N-1)}{2} + N + 2(N-1)$ by collecting the conditions (18), (19), (21) and (22).

We note that the constraints lead to a feasible set of configurations which is sufficiently regular to obtain a well-posed system. More specifically, the set of all overlap free points $\{\boldsymbol{z} \in \mathbb{R}^{6N} \mid \|x_i - x_j\| \geq 2R_i^{\text{hard}} + R_j^{\text{hard}}\}$ is uniformly prox-regular [4], which is the central requirement for existence and uniqueness. For details on the mathematical analysis of inequality-constrained dynamical systems, we refer to [5–7].

##### 2.2.8 Noise

Following [1], we add a small amount of noise which prevents cells from getting locked in an artificial configuration. We note that noise is only added to the cell center positions $x_i$. In terms of the variable $\boldsymbol{z}$, this leads to the following stochastic differential equation

$$\mathrm{d}\boldsymbol{z} = -\nabla_{\boldsymbol{z}} W(\boldsymbol{z})\mathrm{d}t - \sum_{j=1}^{m} \lambda_j \nabla g_j(\boldsymbol{z})\mathrm{d}t + \sigma \mathrm{d}W^t, \tag{23}$$

$$g_j(\boldsymbol{z}) \geq 0, \quad \lambda_j \geq 0 \quad \text{and} \quad \lambda_j g_j(\boldsymbol{z}) = 0 \quad \text{for all } 1 \leq j \leq m \tag{24}$$

where $W^t$ is a $2N$ dimensional Wiener process and the covariance matrix is given by

$$\sigma = \begin{pmatrix} 0 \\ \sigma_x I_{\mathrm{R}^{2N}} \\ 0 \end{pmatrix} \in \mathbb{R}^{6N\times 2N}$$

which effectively adds noise only to the center positions $x_i$ in $\boldsymbol{z} = (a_1, \ldots, x_1, \ldots, b_1, \ldots)$. For mathematical analysis of such systems, we refer to [6].

##### 2.2.9 Initial conditions

For this study, the positions of $N^{\text{init}}$ nuclei are initialized in a domain of size $w^{\text{init}}, h^{\text{init}}$ such that

$$x_i^{\text{init}} \sim \mathcal{U}\left([x^{\text{left}}, x^{\text{right}}] \times [y^{\text{bottom}}, y^{\text{top}}]\right), \tag{25}$$

$$a_i^{\text{init}} = \begin{pmatrix} x^{\text{left}} + \frac{i-1}{N-1} w^{\text{init}} \\ h^{\text{init}} \end{pmatrix}, \tag{26}$$

$$b_i^{\text{init}} = \begin{pmatrix} x^{\text{left}} + \frac{i-1}{N-1} w^{\text{init}} \\ 0 \end{pmatrix} \tag{27}$$

for $1 \leq i \leq N^{\text{init}}$ where

$$x^{\text{right}} = \frac{1}{2} w^{\text{init}}, \quad x^{\text{left}} = -x^{\text{left}}, \tag{28}$$

$$y^{\text{bottom}} = \frac{1}{2} h^{\text{init}}, \quad y^{\text{top}} = \frac{1}{2} h^{\text{init}}. \tag{29}$$

We sort all nuclei positions initially with respect to the $x$-coordinate, e.g. such that $x(x_1^{\text{init}}) \leq x(x_2^{\text{init}}) \leq \cdots \leq x(x_N^{\text{init}})$. We note that the EMT cells will later be located in the middle of the tissue.

For each initial cell, we pick a random cell cycle length according to (3), and then we pick

$$t_i^{\text{birth}} \sim \mathcal{U}([-t_i^{\text{cycle}}, 0]).$$

This approach ensures that the cell cycles are not artificially syncronised initially.

The inital rest length for the apical-nuclei and basal-nuclei springs are set to

$$\eta_i^{\text{ax}} = \eta_i^{\text{bx}} := \eta^{\text{init}}. \tag{30}$$

##### 2.2.10 Summary of the PSE model

The dynamics of the final model are (23) and (24), but with the addition of a discontinuous change during cell division Section 2.2.5 and the updates of rest lengths as in (4), (5) and (12). For simplicity, it is not explicitly denoted, but the number of cells $N$ and the number of constraints $m$ increase after cell division.

A: Loss of apical adhesion

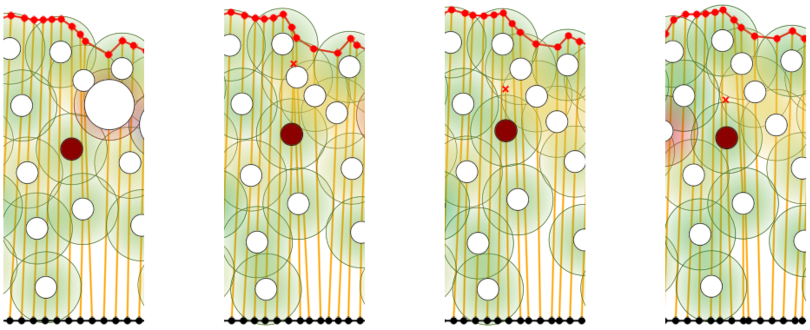


Figure 2: Example for loss of apical adhesion $\boldsymbol{A}$ (cell in red). The apical point (small red cross) first leaves the apical layer and the contraction of the apical-nuclei spring pulls it towards the nuclei.

##### 2.3 The EMT model

In this section, we describe mathematically the rules outlined in Section 1.2.

Let $\mathcal{I}^{\mathrm{EMT}} \subset \mathcal{I} = \{1, \ldots, N\}$ denote the indices of EMT cells. All other cells $\mathcal{I} \setminus \mathcal{I}^{\mathrm{EMT}}$ are control cells which do not perform EMT.

For each EMT cell with index $i \in \mathcal{I}^{\mathrm{EMT}}$ we introduce four extra parameters $T_i^{\mathrm{A}}, T_i^{\mathrm{B}}, T_i^{\mathrm{S}} \in \mathbb{R} \cup \{\infty\}$ and $P_i \in \{0, 1\}$ which define the timings or occurrence of the corresponding EMT-like events $\boldsymbol{A}$, $\boldsymbol{B}$, $\boldsymbol{S}$ and $\boldsymbol{P}$. In the cases, $T_i^{\mathrm{A}} = \infty, T_i^{\mathrm{B}} = \infty$ or $T_i^{\mathrm{S}} = \infty$, the corresponding event will not take place. Similarly, in the case of $P_i = 0$, the cell will not have protrusive activities $\boldsymbol{P}$.

###### 2.3.1 Event *A*: Loss of apical adhesion

At time $t = T_i^{\mathrm{A}}$, we remove the apical point $a_i$ from the apical network and connect the neighbours of the $i$th cell, i.e. we replace the edges $(j_1, i)$ and $(i, j_2)$ with the new edge $(j_1, j_2)$ in $\mathcal{A}$ (which is the set of edges forming the apical layer, see (11)). If the cell is at the boundary, we will only remove the existing apical edge.

Moreover, the desired rest length of the apical-nuclei spring is set to zero, i.e.

$$\eta_i^{\text{ax-des}}(t^+) := 0. \tag{31}$$

The desired apical-nuclei rest length will remain unchanged for the rest of the simulation, in particular, the INM-specific cell event Section 2.2.3 will not apply anymore for the $i$th cell.

##### 2.3.2 Event *B*: Loss of basal adhesion

At time $t = T_i^{\mathrm{B}}$, we remove the basal point $b_i$ from the basal layer and connect the neighbours of the $i$th cell, i.e. we replace the edges $(j_1, i)$ and $(i, j_2)$ with the new edge $(j_1, j_2)$ in $\mathcal{B}$ (see (20)). If the cell is at the boundary, we will only remove the existing basal edge.

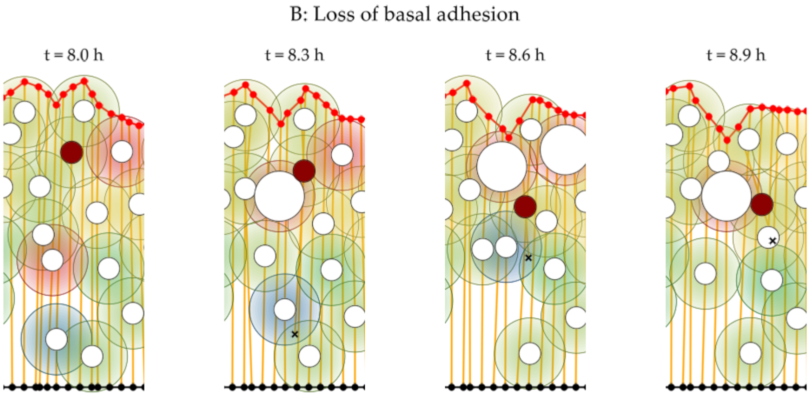


Figure 3: Example for loss of basal adhesion $\boldsymbol{B}$ (cell in red). The basal point (black cross) detaches from the basal layer and the basal-nuclei spring contracts. Notice how the nucleus is pulled towards the basal point during this contraction.

Moreover, we set the desired rest length of the basal-nuclei spring to zero, i.e.

$$\eta_i^{\text{bx-des}}(t^+) := 0. \tag{32}$$

The desired basal-nuclei rest length will remain unchanged for the rest of the simulation, in particular, the INM-specific cell event Section 2.2.3 will not apply anymore for the $i$th cell.

We recall that the constraints for basal points (19), (21) and (22) do only hold for vertices within the basal network. Therefore, after the loss of basal adhesion, the basal point of the $i$th cell can move freely in $\mathbb{R}^2$.

##### 2.3.3 Event *S*: Loss of straightness

At time $t = T_i^{\mathrm{S}}$ we set the stiffness of the bending spring to zero, e.g.

$$\alpha_i^{\text{axb}}(t^+) := 0. \tag{33}$$

As Fig. 4 shows, this increases the ability of the cell to move sideways since the nuclei is not restricted to be along the line that joins the apical and basal sites.

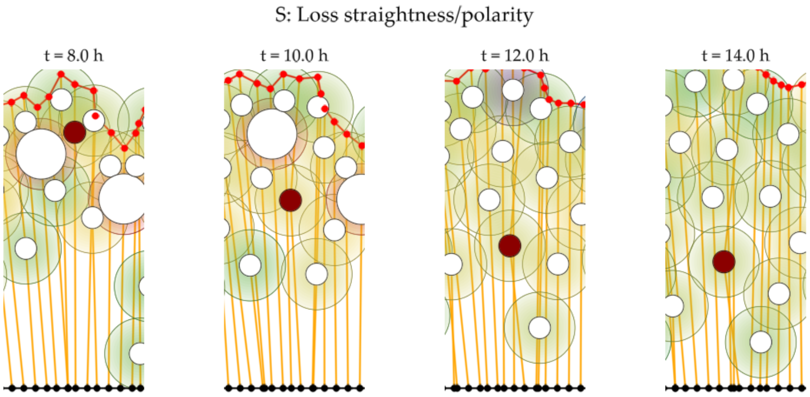


Figure 4: Example for loss of straightness $\boldsymbol{S}$. The EMT cell undergoing event $\boldsymbol{S}$ is in red. Notice how the angle between the apical-nuclei and basal-nuclei springs is no longer straight.

##### 2.3.4 $P$: Protrusive activity

Protrusive activities only occur if $\boldsymbol{B}$ is active and $P_i = 1$. Then, at the time $t = T_i^{\mathrm{B}}$, the $i$th cell will start to self-propel. Cells that do not loss basal adhesion $\boldsymbol{B}$ will never perform protrusive activities $\boldsymbol{P}$.

When the events $\boldsymbol{P}$ starts, we replace the differential equation for the detached basal point $b_i$ (given by (17) and (23)) with the new law

$$\dot{b_i} = \begin{cases} v^{\mathrm{run}} \frac{b_i - x_i}{\|b_i - x_i\|} & \text{if } y(b_i(t)) > -2R^{\mathrm{soft}} \\ 0 & \text{else,} \end{cases} \tag{34}$$

where $y(b_i(t))$ denotes the $y$-coordinate of the basal point. This means that the basal point $b_i$ will move in the direction $b_i - x_i$ at speed $v^{\mathrm{run}}$. The motion stops once the basal point $b_i$ reaches a point that has distance $2R^{\mathrm{soft}}$ from the basal layer (i.e. the full cell must have left the tissue).

The choice of stopping the forward movement if the point is below the basal layer at $+2R^{\mathrm{soft}}$ units of distance is only a convenience, as we do not model the dynamics below the basal layer any further.

We recall that the definition of the basal-nuclei spring (17) leads to the force

$$F^{\mathrm{bx}} = \frac{2\alpha^{\mathrm{bx}}}{R^{\mathrm{soft}} + \eta_i^{\mathrm{bx}}} \left( \|b_i - x_i\| - R^{\mathrm{soft}} - \eta_i^{\mathrm{bx}} \right) \frac{b_i - x_i}{\|b_i - x_i\|} \tag{35}$$

where $\eta_i^{\mathrm{bx}}$ is the rest length of the basal-nuclei spring and $\|b_i - x_i\| - R^{\mathrm{soft}}$ is the current distance between the basal point and the boundary of the nuclei.

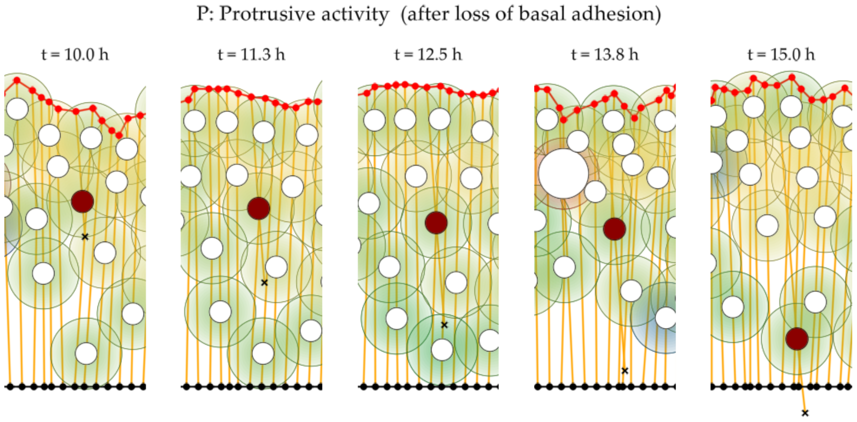


Figure 5: Example of protrusive activities $\boldsymbol{P}$. In this case the cell first lost the basal adhesion at $t = 8\,\mathrm{h}$ (see Fig. 3). The detached basal point is indicated with a black cross and the EMT cell is in red. We note that $\boldsymbol{P}$ is the only EMT event which introduces an active force that leads to extrusion.

As long as the basal point is above the basal layer, this force will be zero since there is nothing the cell can hold onto. But once the basal point is below the basal layer, it will grab onto the basal membrane and the cytoskeleton will pull the cell. We archive this with the following rule

$$\eta_i^{\mathrm{bx}}(t) := \begin{cases} \|x_i - b_i\| - R^{\mathrm{soft}} & \text{if } y(b_i(t)) > 0 \\ 0 & \text{if } y(b_i(t)) \leq 0. \end{cases} \tag{36}$$

This equation replaces (5).

We note, for basal points above the basal layer, we have $F^{\text{basal-cytos}} = 0$, but once $b_i$ is below the basal layer, the cytoskeleton will contract and pull on the nuclei position $x_i$ towards the basal point.

##### 2.3.5 Stop of division

We will not allow EMT cells to further proliferate to simplify statistical evaluations. Therefore, we will use for all EMT cells the cell division rules as in Section 2.2.5 with the additional choice $p^{\text{div-out}} = 100\,\%$. As a result, the EMT cells will effectively reset their cell cycle instead of performing cell division.

### 3 Parameters

#### 3.1 PSE model parameters

The main parameters for the PSE model are given in Tables 1 and 2. These values are equivalent to the choice in [1].

| parameter | value | description |
|---|---|---|
| $N^{\text{init}}$ | 60 | number of cells |
| $h^{\text{init}}$ | 50 µm | initial tissue height |
| $w^{\text{init}}$ | 40 µm | initial tissue width |
| $t^{\text{final}}$ | 54 h | final time |

Table 1: Default model parameters used for simulations. The cell specific parameters are given in Table 2.

#### 3.2 Parameters for EMT model

The only new model parameters compared to the PSE model are the running speed $v^{\text{run}}$ and the timings of EMT-like events.

For simulations, we either randomize the timing for EMT-like events completely (heterogeneous case) or we pick random timings for one, two, or three EMT-like events such that the order of events is the same for all cells (homogeneous case). The corresponding time intervals are given in Table 3. For details on the timings and analysis of the resulting simulations we refer to Section 5.

### 4 Numerical implementation

For the numerical discretisation, we use the position-based dynamics (PBD) method, which is popular in computer graphics [2, 8]. The method uses an explicit Euler step to integrate forces. However, instead of solving the constraints with an implicit solve, the method fuses the first step of a nonlinear projected Gauss-Seidel scheme with the force integration.

We quickly recall the PBD method for a general system in the form

$$\dot{x} = f(x) - \sum_{j=1}^{m} \lambda_j \nabla_x g_j(x) \tag{37}$$

$$g_j(x) \geq 0, \quad \lambda_j \geq 0, \quad g_j(x)\lambda_j = 0. \tag{38}$$

| parameter | value | description |
|---|---|---|
| $R^{\text{soft}}$ | $5\,\mu\text{m}$ | radius of soft repulsion |
| $R^{\text{hard}}$ | $1.66\,\mu\text{m}$ | radius of non-overlap (during S/passive G2 phase) |
| $R^{\text{hard,G2}}$ | $0.7R^{\text{soft}}$ | radius of non-overlap (during active G2, mitosis phase) |
| $\eta^{\text{init}}$ | $7.5\,\mu\text{m}$ | initial rest length of linear springs (cytoskeleton) |
| $\sigma_x^2$ | $0.01\,\mu\text{m}^2\,\text{h}^{-1}$ | diffusion coefficient for nuclei positions |
| $T^{\text{G2}}$ | $30\,\text{min}$ | duration of G2 phase |
| $T^{\text{mitsis}}$ | $30\,\text{min}$ | duration of mitosis |
| $T^{\text{min-cycle}}$ | $10\,\text{h}$ | minimal duration of full cell cycle |
| $T^{\text{max-cycle}}$ | $21\,\text{h}$ | maximal duration of full cell cycle |
| $k^{\text{aa}}$ | $1\,\text{h}^{-1}$ | adaption speed of apical-apical spring |
| $k^{\text{ax}}$ | $5\,\text{h}^{-1}$ | adaption speed of apical-nuclei spring |
| $k^{\text{bx}}$ | $5\,\text{h}^{-1}$ | adaption speed of basal-nuclei spring |
| $\alpha^{\text{aa}}$ | 5 | stiffness of apical-apical spring |
| $\alpha^{\text{ax}}$ | 2 | stiffness of apical-nuclei spring |
| $\alpha^{\text{bx}}$ | 2 | stiffness of basal-nuclei spring |
| $\alpha^{\text{xx}}$ | 1 | stiffness of nuclei-nuclei repulsion |
| $\alpha^{\text{axb}}$ | 15 | stiffness bending spring (straightness) |

Table 2: Default cell parameters for the PSE model. We refer to [1] for details on the calibration of the model.

where $f$ denotes the forces and $g_1, \ldots, g_m : \mathbb{R}^d \to \mathbb{R}$ denote inequality constraint functions with their corresponding multipliers $\lambda_1, \ldots, \lambda_m$.

For the formulation of numerical methods, it is useful to define the corresponding feasible sets

$$S_j := \{x \in \mathbb{R}^d \mid g_j(x) \geq 0\} \quad \text{and} \quad S = \bigcap_{j=1}^{m} S_j. \tag{39}$$

We can equivalently write the dynamics (37) and (38) using the set $S$ as it is explained in [9].

The position-based dynamics method applied to this first-order system reads

$$x_{k+1} = P_{S_m} \circ \cdots \circ P_{S_1}(x_k + \Delta t f(x_k)). \tag{40}$$

For a convergence proof of the PBD method in this context, we refer to [3].

Applied to our context, the constraints are the functions defined Section 2.2.7, which implement the non-overlap between hard spheres and the dynamics of the basal points.

| parameter | value | description |
|---|---|---|
| $N^{\mathrm{EMT}}$ | 1 or 11 | number of EMT cells |
| $v^{\mathrm{run}}$ | $1\,\mu\mathrm{m}\,\mathrm{h}^{-1}$ | running speed during P |
| $[T^{\min}, T^{\max}]$ | $[6\,\mathrm{h}, 24\,\mathrm{h}]$ | interval for EMT-like events (heterogeneous case) |
| $[T^{1,1}, T^{1,2}]$ | $[6\,\mathrm{h}, 24\,\mathrm{h}]$ | interval of single EMT-like event |
| $[T^{2,1}, T^{2,2}]$ | $[6\,\mathrm{h}, 15\,\mathrm{h}]$ | first interval for two EMT-like events |
| $[T^{2,2}, T^{2,3}]$ | $[15\,\mathrm{h}, 24\,\mathrm{h}]$ | second interval for two EMT-like events |
| $[T^{3,1}, T^{3,2}]$ | $[6\,\mathrm{h}, 12\,\mathrm{h}]$ | first interval for three EMT-like events |
| $[T^{3,2}, T^{3,3}]$ | $[12\,\mathrm{h}, 18\,\mathrm{h}]$ | second interval for three EMT-like events |
| $[T^{3,3}, T^{3,4}]$ | $[18\,\mathrm{h}, 24\,\mathrm{h}]$ | third interval for three EMT-like events |

Table 3: Interval of opportunity for simulations with one, two, or three EMT-like events. The concrete values per cell are uniformly distributed within these intervals. See Sections 5.3 and 5.4 for further details on the random parameters.

In this case, all projections $P_{S_j}$ are explicitly known and easy to compute. For example, let us use $S_{ij}$ for the sets of all pairwise non-overlap between the hard spheres of the cells, which is the feasible set corresponding to the inequality constraint (18) which we might denote as

$$g_{ij}(\boldsymbol{z}) = \|x_i - x_j\| - R_i^{\mathrm{hard}} - R_j^{\mathrm{hard}} \geq 0.$$

For a pair of cells, the projection is given by

$$P_{S_{ij}}(\boldsymbol{z}) = \begin{cases} \boldsymbol{z} - \nabla_{\boldsymbol{z}} g_{ij}(\boldsymbol{z}) \frac{g_{ij}(\boldsymbol{z})}{\|\nabla_{\boldsymbol{z}} g_{ij}(\boldsymbol{z})\|^2} & \text{if } g_{ij}(\boldsymbol{z}) < 0, \\ \boldsymbol{z} & \text{else,} \end{cases}$$

which simply shifted the two positions $x_i, x_j$ sufficiently apart from each other.

With the constraints as in Section 2.2.7 denoted as $S_1, \ldots, S_m$ and for a given initial value $\boldsymbol{z}_0 \in \mathbb{R}^{6N}$, one time-step of the position based dynamics method reads

$$\Phi^{\mathrm{PBD}}(\boldsymbol{z}_k) = P_{S_m} \circ \cdots P_{S_1}(\boldsymbol{z}_k - \Delta t \nabla_{\boldsymbol{z}} W(\boldsymbol{z}_k)).$$

In addition to the integration of the mechanical system, we also need to update parameters, perform EMT events and add noise. Since the PBD method benefits from smaller time steps without external interruptions, the parameter $n^{\text{substeps}}$ determines the number of steps of the PBD method until phenotype updates and noise are added. The resulting method reads

$$\boldsymbol{z}_{k+1} = \left(\Phi^{\mathrm{PBD}}(\boldsymbol{z}_k)\right)^{n^{\text{substeps}}} + \sigma\sqrt{\Delta t} W_k, \tag{41}$$

$$p_{k+1} = \text{cell events}(p_k), \tag{42}$$

where $W_k$ are i.i.d. normally distributed samples of dimension $2N$ and $p_k, p_{k=1}$ denote the cell parameters which do not follow an differential equations, such as the desired rest lengths $\eta^{\mathrm{ax-des}}, \eta^{\mathrm{bx-des}}$ and other quantities like the number of cells $N$ or the hard sphere radii $R_i^{\mathrm{hard}}$.

The used numerical parameters are given in Table 4.

Compared to the quasi-steady state model in [1], this new formulation of the PSE model combined with the PBD method led to a considerable speedup and improved the numerical stability. However, the simulation results are, in practical terms, identical.

| parameter | value | description |
|---|---|---|
| $\Delta t$ | $0.01\,\mathrm{h}$ | time step (between cell events) |
| $n^{\mathrm{substeps}}$ | 40 | number of substeps of the PBD method |

Table 4: Numerical parameters used for simulations.

### 5 Extrusion metrics and analysis of ensemble simulations

#### 5.1 Notation

In the following, we will consider ensemble simulations of the model, which generate $n^{\mathrm{rep}} \in \mathbb{N}$ sample trajectories $\boldsymbol{z}_j : [0, t^{\mathrm{final}}] \to \mathbb{R}^{6N}$ for $1 \leq j \leq n^{\mathrm{rep}}$. We refer to the indices of EMT cells as $\mathcal{I}^{\mathrm{EMT}}$, and the control cells have labels $\mathcal{I}^{\mathrm{control}}$.

We also recall that we refer to $y$ coordinates of a position $x_i \in \mathbb{R}^2$ as $y(x_i)$.

Finally, we denote the line connecting two points $\boldsymbol{a}, \boldsymbol{b} \in \mathbb{R}^2$ as

$$\overline{a\,b} := \{\lambda a + (1-\lambda)b \mid \lambda \in [0,1]\}. \tag{43}$$

#### 5.2 Definition of apical and basal extrusion and basal positioning

The three most important quantities for our analysis of EMT are the numbers of *apically extruded cells*, *basally extruded cells* and the number of EMT cells *below control cells*.

To quantify the position of a cell within the tissue, we introduce the *apical-basal scale (AB scale)*, which is zero at the basal layer and one at the apical layer. It allows us to compare the relative positions of cell nuclei independent of the current development stage.

In the EMT model, not all cells have their apical point on the apical layer. To define the AB scale, we will therefore use the projection of the nuclei onto the apical layer as a reference point. For a fixed state $\boldsymbol{z} = (a_1, \ldots, x_1, \ldots, b_1, \ldots) \in \mathbb{R}^{6N}$ with apical network $\mathcal{A}$ we define the apical layer $L_{\boldsymbol{z}}$ as the path of lines that passes through all points $a_i$

forming the apical surface, i.e.

$$L_{\boldsymbol{z}} := \bigcup_{(i,j)\in\mathcal{A}} \overline{a_i\, a_j}. \tag{44}$$

For the $i$th nucleus at position $x_i \in \mathbb{R}^2$, we can compute the closest point on the apical layer as

$$A_i^{\mathrm{ref}}(\boldsymbol{z}) := P_{L_{\boldsymbol{z}}}(x_i)$$

where P denotes the projection operator onto $L_{\boldsymbol{z}}$. Since the basal layer is fixed at the $x$-axis, we can define the *position of a cell on the AB scale* as

$$y_i^{\mathrm{AB}}(\boldsymbol{z}) := \frac{y(x_i)}{y(A_i^{\mathrm{ref}}(\boldsymbol{z}))}.$$

To denote if cells are apically or basally extruded, we define the following indicator functions:

$$\delta_i^{\mathrm{basal-extr}}(\boldsymbol{z}) := \begin{cases} 1 & \text{if } y_i^{\mathrm{AB}}(\boldsymbol{z}) < 0, \\ 0 & \text{else,} \end{cases} \qquad \delta_i^{\mathrm{apical-extr}}(\boldsymbol{z}) := \begin{cases} 1 & \text{if } y_i^{\mathrm{AB}}(\boldsymbol{z}) > 1, \\ 0 & \text{else.} \end{cases}$$

Moreover, we say a cell is in below the control cells if its nucleus is below all control cells with respect to the apical-basal scale. Hence, we define

$$\delta_i^{\mathrm{below-control}}(\boldsymbol{z}) := \begin{cases} 1 & \text{if } y_i^{\mathrm{AB}}(\boldsymbol{z}) \le \min_{j\in\mathcal{I}^{\mathrm{control}}} y^{\mathrm{AB}}(x_j), \\ 0 & \text{else.} \end{cases} \tag{45}$$

#### 5.3 Homogeneous case and estimated probability of extrusion

In the homogeneous case, all EMT cells in the tissue perform the same EMT-like events. Figure 2 to Figure 4 in the main document are based on simulations, where the selection and order of EMT-like events is the same for all EMT cells. The time intervals of opportunity are given in Table 3.

For example, for the scenario $\boldsymbol{ABS}$, the timings for the second EMT-like event $\boldsymbol{B}$ are uniformly distributed in $[T^{3,2}, T^{3,3}]$, e.g.,

$$T_i^B \sim \mathcal{U}(T^{3,2}, T^{3,3}) \quad \text{i.i.d. for } i \in \mathcal{I}^{\mathrm{EMT}}, \tag{46}$$

where $\mathcal{U}$ denotes the uniform distribution on the time interval.

We use this randomized choice of EMT timings for ensemble simulations with $n^{\mathrm{rep}}$ sample trajectories denoted as $(\boldsymbol{z}_j)_j$ for $1 \le j \le n^{\mathrm{rep}}$. Based on these samples, we estimate the probability of apical extrusion for given scenarios as

$$p^{\mathrm{apical-extr}}(t) := \frac{1}{n^{\mathrm{rep}}|\mathcal{I}^{\mathrm{EMT}}|} \sum_{j=1}^{n^{\mathrm{rep}}} \sum_{i\in\mathcal{I}^{\mathrm{EMT}}} \delta_i^{\mathrm{apical-extr}}(\boldsymbol{z}_j(t)). \tag{47}$$

Equivalently, we define $p^{\mathrm{basal-extr}}$ and $p^{\mathrm{below-control}}$. These values are plotted in Figures 2 to 4.

For the homogeneous, case we used $n^{\mathrm{rep}} = 1000$ to obtain a sufficiently accurate statistical estimation.

#### 5.4 Heterogeneous case

In Figure 5 and Figure 6 of the main document, we not only randomize the timings of EMT-like events, but also allow EMT cells in one simulation to follow different scenarios, that is, a different selection and order of EMT-like events. Moreover, we also introduce heterogeneity with respect to interkinetic nuclear migration (INM) and protrusive activities $\boldsymbol{P}$ by restricting these events to a subset of the EMT cells.

More specifically, we consider the case that each EMT cell performs INM (Section 2.2.3) with a likelihood of $50\%$.

Similarly, we also consider the case that each individual EMT cell has a probability of $50\%$ for protrusive activities after loss of basal adhesion $\boldsymbol{B}$.

We randomize the selection and order of EMT-like events, by selecting the event times $T_i^{\mathrm{A}}, T_i^{\mathrm{B}}$ and $T_i^{\mathrm{S}}$ for $i \in \mathcal{I}^{\mathrm{EMT}}$ as i.i.d. random values with distribution

$$\begin{aligned} &\mathbb{P}[T_i^* = \infty] = 0.3 \quad \text{and} \\ &\mathbb{P}[t < T_i^* < s] = 0.7 \frac{s-t}{T^{\max} - T^{\min}} \quad \text{for all } T^{\min} \le t \le s \le T^{\max}, \end{aligned}$$

where $*$ is a placeholder for $\mathrm{A}, \mathrm{B}, \mathrm{S}$.

The extrusion probabilities are then estimated as in (47). We note that due to the larger variability, we require more samples than in the homogeneous case. The results presented in the main document use data from $n^{\mathrm{rep}} = 150.000$ simulations.

#### 5.5 Correlation analysis

The correlation factors presented in Figure 6 of the main document are computed as follows.

Using the heterogeneous setup from Section 5.4 with $N^{\mathrm{EMT}} = 11$, we investigate the correlation between the randomized parameters and the resulting extrusion statistics.

For the correlation analysis, we define the first and last event time as

$$\begin{aligned} T_i^{\mathrm{first}} &:= \min(T_i^A, T_i^B, T_i^S), \\ T_i^{\mathrm{last}} &:= \max(T_i^A, T_i^B, T_i^S) \qquad \text{(ignoring cases of } T_i^* = \infty) \end{aligned}$$

and we introduce the binary variables

$$A_i = \begin{cases} 1 & \text{if } T_i^A < \infty, \\ 0 & \text{else,} \end{cases} \quad B_i = \begin{cases} 1 & \text{if } T_i^B < \infty, \\ 0 & \text{else,} \end{cases} \quad S_i = \begin{cases} 1 & \text{if } T_i^S < \infty, \\ 0 & \text{else.} \end{cases}$$

Moreover, we set $P_i = 1$ if $\boldsymbol{P}$ is potentially possible for a cell and $P_i = 0$ otherwise.

We also consider the position $i \in \mathcal{I}^{\mathrm{EMT}}$ within the EMT cluster as a random choice. We denote this random index as $\alpha \sim \mathcal{U}(\mathcal{I}^{\mathrm{EMT}})$.

Based on these notations, we define the following random variables

$$\begin{aligned} A &= A_\alpha, \quad B = B_\alpha, \quad P = P_\alpha, \quad S = S_\alpha, \\ t_* &= T_\alpha^* \quad \text{for } * \in \{A, B, P, S\}, \\ \Delta t_{\mathrm{emt}} &= T_\alpha^{\mathrm{last}} - T_\alpha^{\mathrm{first}}, \\ y_{\mathrm{init}} &= y(x_\alpha^{\mathrm{init}}), \\ y_{\mathrm{emt}} &= y(x_\alpha(T_\alpha^{\mathrm{first}})), \\ y_* &= y(x_\alpha(T_\alpha^*)) \quad \text{for } * \in \{A, B, P, S\}. \end{aligned}$$

Using the sample trajectories $(\boldsymbol{z}_j)_j$ from the ensemble simulations, we compute the correlation between these random variables and $\delta^{\mathrm{apical-extr}}(t^{\mathrm{final}})$ and $\delta^{\mathrm{basal-extr}}(t^{\mathrm{final}})$ are presented in Figure 6 of the main document. We note that not all variables are always well-defined. In these cases we filtered samples with undefined values for the estimation of the correlation. Specifically, we removed all cases with no or only one EMT-like event for the correlation with $\Delta t_{\mathrm{emt}}$. We removed cases with no EMT-like event when computing correlation with $y_{\mathrm{emt}}$ and finally, we considered only data for $t_*$ and $y_*$ whenever the corresponding EMT-like event was active.
