## Supplementary material for "Modelling variability and heterogeneity of EMT scenarios highlights nuclear positioning and protrusions as main drivers of extrusion": Supp Movies Legends

**Supplementary Movie Legends**

**Movie S1**

Examples of simulations showing the impact of loss of cell-cell adhesion (***A***) in single EMT-like cells with interkinetic movements (top panels) or in a group of EMT-like cells with interkinetic movements (bottom panel). All epithelia are oriented apical side up. EMT-like cells are in red. Cells in mitosis are displayed with an enlarged nucleus. Related to Figure 2.

**Movie S2**

Examples of simulations showing the impact of loss of anchorage to basal domain (***B***) in single EMT-like cells with interkinetic movements (top panels) or in a group of EMT-like cells with interkinetic movements (bottom panel). All epithelia are oriented apical side up. EMT-like cells are in red. Cells in mitosis are displayed with an enlarged nucleus. Related to Figure 2.

**Movie S3**

Examples of simulations showing the impact of loss of anchorage to basal domain (***B***) in single EMT-like cells (top panels) or in a group of EMT-like cells (bottom panels) with interkinetic movements (left panels) or without INM (right panels). All epithelia are oriented apical side up. EMT-like cells are in red. Cells in mitosis are displayed with an enlarged nucleus. Related to Figure 2.

**Movie S4**

Examples of simulations showing the impact of a scenario with three EMT-like events (***ABS***, loss of cell-cell adhesion followed by loss of anchorage to the basal domain and relaxation of the straightness factor) in single EMT-like cells (top panels) or in a group of EMT-like cells (bottom panels) without interkinetic movements, without protrusions (***P***, left panels) or with protrusions (right panels). All epithelia are oriented apical side up. EMT-like cells are in red. Cells in mitosis are displayed with an enlarged nucleus. Related to Figure 2.
